# Transcription of a centromere-enriched retroelement and local retention of its RNA are significant features of the CENP-A chromatin landscape

**DOI:** 10.1101/2024.01.14.574223

**Authors:** B Santinello, R Sun, A Amjad, SJ Hoyt, L Ouyang, C Courret, R Drennan, L Leo, AM Larracuente, L Core, RJ O’Neill, BG Mellone

**Affiliations:** Department of Molecular and Cell Biology, University of Connecticut, Storrs, CT, US; Department of Biology, University of Rochester, Rochester, NY, US; Dipartimento di Biologia e Biotecnologie Charles Darwin, Sapienza University of Rome, 00185 Rome, Italy; Present address: RNA editing Lab, Onco-Haematology Department, Genetics and Epigenetics of Pediatric Cancers, Bambino Gesù Children Hospital, IRCCS, 00146 Rome, Italy; Institute for Systems Genomics, University of Connecticut, Storrs CT, US; 6Department of Genetics and Genome Sciences, UConn Health, Farmington, CT, US

## Abstract

Centromeres depend on chromatin containing the conserved histone H3 variant CENP-A for function and inheritance, while the role of centromeric DNA repeats remains unclear. Retroelements are prevalent at centromeres across taxa and represent a potential mechanism for promoting transcription to aid in CENP-A incorporation or for generating RNA transcripts to maintain centromere integrity. Here, we probe into the transcription and RNA localization of the centromere-enriched retroelement *G2/Jockey-3* (hereafter referred to as *Jockey-3*) in *Drosophila melanogaster*, currently the only *in vivo* model with assembled centromeres. We find that *Jockey-3* is a major component of the centromeric transcriptome and produces RNAs that localize to centromeres in metaphase. Leveraging the polymorphism of *Jockey-3* and a *de novo* centromere system, we show that these RNAs remain associated with their cognate DNA sequences in *cis*, suggesting they are unlikely to perform a sequence-specific function at all centromeres. We show that *Jockey-3* transcription is positively correlated with the presence of CENP-A, and that recent *Jockey-3* transposition events have occurred preferentially at CENP-A-containing chromatin. We propose that *Jockey-3* contributes to the epigenetic maintenance of centromeres by promoting chromatin transcription, while inserting preferentially within these regions, selfishly ensuring its continued expression and transmission. Given the conservation of retroelements as centromere components through evolution, our findings have broad implications in understanding this association in other species.

## Introduction

Genome partitioning during cell division is dependent on specialized chromosomal structures known as centromeres, which mediate kinetochore assembly. This process is crucial for establishing robust connections between chromosomes and spindle microtubules, essential for the precise segregation of chromosomes. Centromeric chromatin is marked by the presence of nucleosomes containing the histone H3 variant CENP-A (also known as Cid *Drosophila*)(1, 2), which initiates the recruitment of additional centromeric and kinetochore proteins (3). Centromeres are paradoxical in that they play a highly conserved function across eukaryotes yet are amongst the most rapidly evolving regions of genomes. Centromeres are also dynamic – they can reposition in individuals (neocentromeres)(4) and become fixed in a population (evolutionary new centromeres)(5). Despite being able to reposition, centromeres are typically associated with large highly repetitive sequences whose role in 880pcentromere identity remains elusive.

Transcripts emanating from centromeres have been observed in a myriad of systems, including budding yeast (6, 7), human cells (8–11), frog egg extracts (12, 13), maize (14), and marsupials (15). Transcription at centromeres has been shown to be coupled to *de novo* centromere formation (16) and neocentromere formation in humans (17–19). In addition, centromeric transcription is critical for programmed histone exchange in *S. pombe* (20), for the stabilization of newly formed CENP-A nucleosomes in *Drosophila* cells (21), and for Human Artificial Chromosome formation (22). These studies suggest that centromeric DNA may contribute to centromere identity through its ability to be transcribed. Other studies have also implicated a role for centromere-derived transcripts as noncoding RNAs important for centromere integrity (8, 9, 12, 14, 15). Indeed, in some cases centromeric transcripts have been detected associated with centromeric proteins (9, 12, 13), suggesting a role beyond being a byproduct of transcription. However, whether the interaction with centromere proteins is sequence-specific remains unresolved. Furthermore, both the functional impact of these RNAs, as well as the extent of their prevalence across different systems, are still not fully understood.

Consistent with the existence of centromeric transcripts, elongating RNA polymerase II accumulates at mitotic centromeres in *Drosophila* S2 cells (21, 23) and nascent transcription can be detected at the centromere of *Drosophila* S2 cells in mitosis and G1 (21). However, the RNA products of such centromeric transcription in *Drosophila* are unknown. A previous study analyzed the role of a non-coding RNA produced by a satellite of the 1.688 family, showing that its depletion affects accurate chromosome segregation and centromere integrity (23). However, the largest block of this satellite is located within pericentric heterochromatin on the X (24) and its RNA product does not localize to centromeres (21). Therefore, its contributions to centromere segregation accuracy might be unrelated to centromeric defects.

The centromeres of *Drosophila melanogaster* have been recently annotated (24), providing a unique opportunity to directly analyze transcripts associated with centromeres. *Drosophila* has five chromosomes (X; Y; 2; 3; and 4), each harboring a unique centromere differing in repeat composition and organization. The centromeres are composed of islands of complex repeats enriched in retroelements embedded in large arrays of simple satellites. CENP-A occupies primarily the islands, which are between 101-171-kb, extending only partially to the flanking satellites. All of the repeats present at *Drosophila* centromeres are also present elsewhere in the genome, yet a subset of retroelements are enriched at centromeres (24). Only one element, the non-LTR retroelement *G2/Jockey-3* (henceforth *Jockey-3*), is shared between all centromeres and is conserved at the centromeres of *D. simulans*, a species that diverged from *D. melanogaster* 2.5 million years ago (25) and that displays highly divergent centromeric satellites (26, 27). Retroelements are conserved centromere-associated elements across taxa. In plants, these elements have been proposed to help maintain centromere size and increase the repeat content of neocentromeres (28). Additionally, retroelements could contribute to centromere function in two ways: either by facilitating localized transcription thought to promote CENP-A incorporation (16, 21, 29–33) or by generating transcripts with non-coding roles in maintaining centromere integrity as postulated for other repeats (9, 12, 13, 34). Whether retroelements play such roles remains unknown.

Here, we investigate the expression and RNA localization of the conserved centromere-enriched retroelement *Jockey-3*. Nascent transcription profiling and total RNA-seq in *Drosophila* embryos show that centromeric and non-centromeric copies of *Jockey-3* are actively transcribed. Using single-molecule RNA FISH combined with immunofluorescence for the centromere protein CENP-C, we show that, during mitosis, *Jockey-3* RNA transcripts localize primarily to centromeres and remain associated with their locus of origin in *cis*. We also show that the presence of CENP-A chromatin is strongly correlated with transcription at both centromeric and non-centromeric full-length *Jockey-3* copies. Furthermore, we find that recent *Jockey-3* transposition events occur preferentially at CENP-A containing domains across the genome. *De novo* centromere formation *in vivo* using a LacI/lacO tethering system results in the accumulation of lacO transcripts at the *de novo* centromere in mitosis, suggesting that even in the absence of *Jockey-3* or any other centromere-enriched repeats, CENP-A chromatin formation is coupled with transcription *in vivo*. Our work supports a model whereby the *Jockey-3* retroelement targets CENP-A chromatin for its selfish propagation while contributing to CENP-A maintenance through transcription. CENP-A chromatin in itself promotes transcription when artificially assembled. This work provides a framework to understand the persistent association between retroelements and centromeres through evolution.

## Results

### The transcriptional profile of *Drosophila* centromeres

Transcription of centromeric DNA has been implicated in centromere maintenance in both a sequence-independent manner and through the action of specific transcripts (33, 35–37). In *Drosophila*, only a few known satellite transcripts have been identified (21, 23, 38, 39), but these are either pericentric or not derived from the sequences most highly associated with CENP-A (24). The availability of annotated centromeres for the *Drosophila* laboratory strain iso-1 and the discovery that these centromeres contain retroelements (24) present a unique opportunity to examine transcription across these previously unresolved regions of the genome and explore the correlation with CENP-A occupancy. To identify nascent transcripts, we generated libraries for Precision Nuclear Run-On sequencing (PRO-seq), which detects nascent transcription from RNA polymerase with nucleotide resolution (40) from 0-12h old embryos and 3rd instar larval brains. We also generated RNA-seq libraries for the same type of samples, providing a catalog of stable transcripts. Plotting our PRO-seq data for all genes showed the expected transcriptional profile with a peak at the 5 ’of genes, confirming successful capture of elongating RNA polymerase (**Fig. S1**). Since none of the repeats found at the centromeres are unique to these regions and PRO-seq and RNA-seq generate short-read data, nascent transcripts identified by PRO-seq did not map uniquely to the centromeres using standard mapping methods. To overcome this limitation and determine if any nascent transcripts emanate from centromeric sequences, we adapted a mapping-dependent method recently developed for the human repeats transcriptome (11) to our *Drosophila* datasets. For each dataset, Bowtie 2 default “best match” reports a single alignment for each read providing locus-level transcription profiles (lower bounds); unfiltered Bowtie/Bowtie 2 k-100 mapping reports up to 100 mapped loci for each read, providing over-fitted and locus-level transcriptional profiles (upper bounds); and single copy k-mer filtering, with 21-mers for PRO-seq and 51-mers for RNA-seq data applied to Bowtie k-100, reveals the intermediate bounds of locus-level transcription (**Fig. 1A**). This k-mer filtering requires a given read alignment to overlap with an entire single copy k-mer in the assembly in order to be retained. Together, these different approaches provide a more complete representation of the true transcriptional landscape of centromeres.

**Figure 1.**
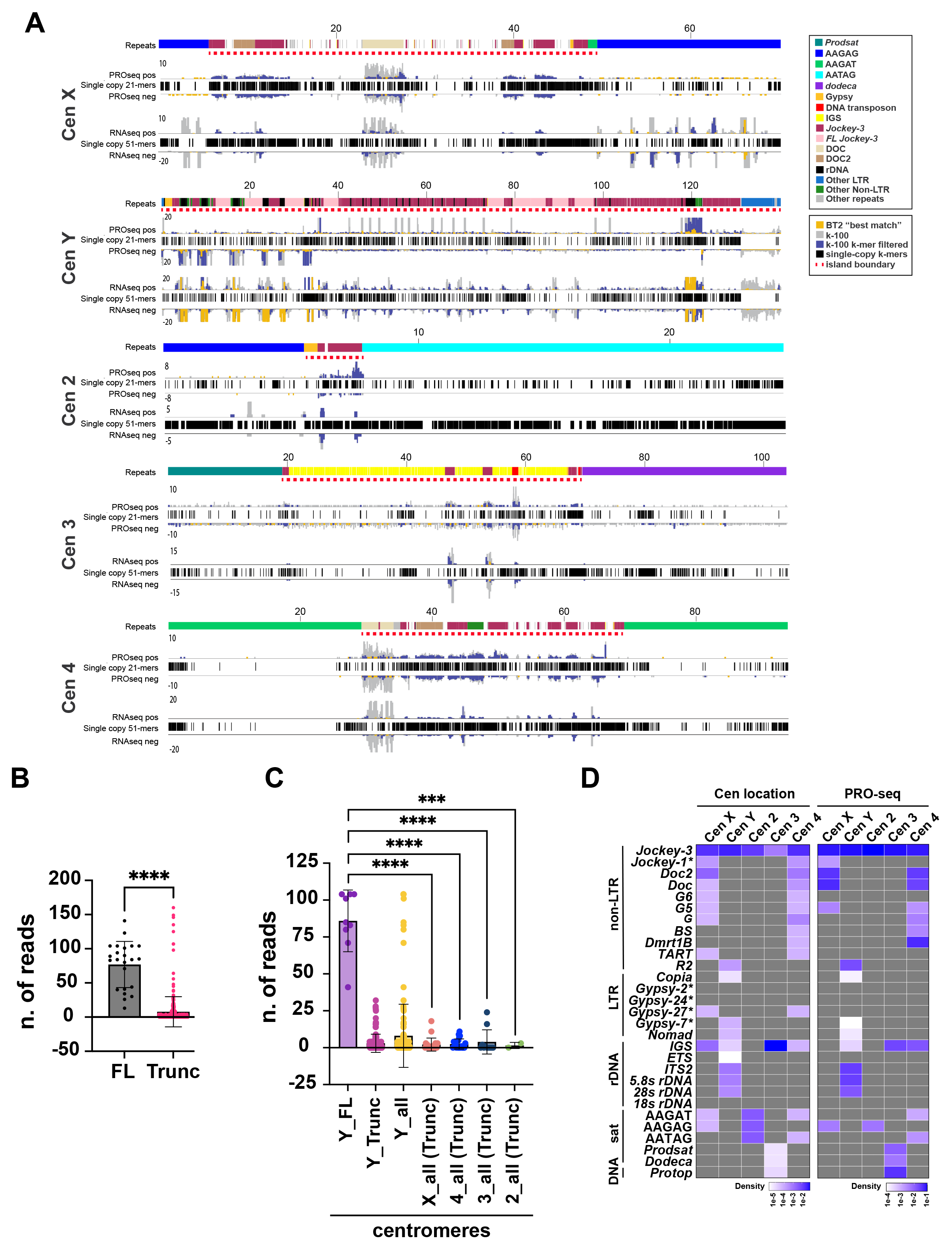
The transcriptional profile of Drosophila centromeres reveals Jockey-3 as a major transcribed element. **A** PRO-seq, RNA-seq signals for 0-12h embryos across all *D. melanogaster* centromeres. Top track shows sense, bottom, antisense. Tracks show read coverage with three mapping methods: Bowtie 2 default “best match” (“lower bounds”; yellow), over-fit (“upper bounds”; gray) and a filtered over-fit (“intermediate bounds”; blue). For PRO-seq we used Bowtie k-100 for over-fit, and Bowtie k-100 unique 21-mer filtered for intermediate bounds. For RNA-seq we used Bowtie2 k-100 for over-fit and Bowtie2 k-100 unique 51-mer filtered for intermediate bounds. Repeat annotation is shown on top (see legend for details), with unique 21 and 51-mers (black) used for the filtering shown below. The k-mer tracks illustrate the regions that lack sequence specificity and are therefore most prone to read loss through k-mer filtering. Coordinates shown are kilobases. The boundaries of centromere islands are demarcated by a red dashed line. **B** PRO-seq read density scatter boxplot comparison between full-length and truncated (minus three outliers) *Jockey-3* copies, regardless of genome location. Mapping was done with Bowtie 2 default “best match” using paired-end reads, post-deduplication. An unpaired t-test determined a statistically significant difference (****; p < 0.0001; Student’s t-test). Standard deviation error bars are shown. **C** PRO-seq read density scatter-boxplot comparisons of centromeric *Jockey-3* copies split by chromosome and whether they are full-length vs. truncated. Since chromosome Y includes both full-length and truncated copies, a third bar was included encompassing all copies; all three bars are indicated by a dashed box. Mapping was performed with Bowtie 2 default “best match” using paired-end reads, post-deduplication. FL, full-length, Trunc, truncated. Note that only the Y centromere contains FL copies, hence for all other centromeres ‘All’ is made up of only truncated copies. An unpaired t-test determined a statistically significant difference (****; p < 0.0001; ***, p < 0.001; Student’s t-test). All other comparisons with Y_FL have p < 0.0001 (omitted in plot). Error bars show the standard deviation. **D** Left, density plot of all repetitive elements on each candidate centromere contig grouped by type as in Chang et al (non-LTR retroelements, LTR retroelements, rDNA-related sequences, simple satellites, and DNA transposon) using an updated genome annotation from Hemmer et al (61). An * indicates annotations based on similarity to retroelements in other *Drosophila* species: *Jockey-1* and *Gypsy-2* are from *D. simulans*, *Gypsy-24* and *Gypsy-27* are from *D. yakuba*, and *Gypsy-7* is from *D. sechellia*. Right, density plots showing PRO-seq reads (k-100 filtered) for a given repeat (see label from C) normalized by the total number of reads mapping to each contig. Density scale is shown in blue. Gray indicates zero copies/reads for a given repeat.

We observe nascent transcription at all centromeres, particularly within the islands (**Fig. 1A**). Based on our statistical tests, *Jockey-3* nascent transcripts emerge primarily from full-length *Jockey-3* elements (**Fig. 1B, Fig. S2; Table S1**), 9/23 of which are within the Y centromere, while the rest (14/23) are non-centromeric (**Table 1**). Both centromeric and non-centromeric truncated *Jockey-3* elements are transcribed (**Table S1**), suggesting that the putative promoter at the 5 ’end (Hemmer et al., 2023) is not required for *Jockey-3* transcription. When we compared the number of *Jockey-3* reads mapping to each of the centromeres, classified based on whether they are full-length or truncated, we observed significantly more reads coming from full-length *Jockey-3* insertions within the Y centromere compared to all others (**Fig. 1C**).

**Table 1:** Summary of the location of truncated and full-length (FL) *Jockey-3* insertions with estimated age. Table showing the distribution of *Jockey-3* copies per centromere, across all centromeres, and across non-centromeric loci. The number of copies with <1% divergence from the *Jockey-3* consensus were deemed ‘young’ and are indicated in parenthesis. The difference between ‘young’ and total corresponds to the number of ‘old’ copies (≥1% divergence).

| CEN | All | CEN X | CEN Y | CEN 2 | CEN 3 | CEN 4 | CEN (all) | nonCEN |
| --- | --- | --- | --- | --- | --- | --- | --- | --- |
| **All *Jockey-3*** | 329 | 21 | 147 | 2 | 11 | 21 | 202 | 127 |
| **FL *Jockey-3*** | 23 (16) | 0 (0) | 9 (4) | 0 (0) | 0 (0) | 0 (0) | 9 (4) | 14 (12) |
| **Truncated *Jockey-3*** | 306 (26) | 21 (3) | 138 (5) | 2 (0) | 11 (0) | 21 (1) | 193 (9) | 113 (17) |

Similarly to nascent RNA data, RNA-seq profiles from embryos reveal the presence of transcripts predominantly mapping to the islands, with low levels of satellite transcripts, with the notable exception of AAGAG on the X centromere, which shows more expression in this dataset (**Fig. 1A**). PRO-seq from larval brains (**Fig. S3**), as well as from 0-4h and 4-8h old embryos (data not shown) also showed very similar transcriptional profiles. In contrast, RNA-seq profiles from larval brains showed more transcripts mapping to flanking satellites compared to what we observed in the embryos datasets (**Fig. S3**).

To determine more quantitatively which centromere-associated repeats are transcribed, we generated read count plots for each of the repeats found within the centromere contigs. We recreated a density plot of all repetitive elements as in (24) using an updated genome annotation (41) to show how many copies of each repeat are present within each of the centromere contigs (**Fig. 1D**-left plot). We then generated a density heat map for the PRO-seq 0-12h embryos dataset, which displays the total read count for each repeat normalized by the total reads mapping to that contig. This heat map shows that *Jockey-3* is highly expressed at all centromeres relative to other centromeric repeats (**Fig. 1D**-right plot **and Table S2**). Several repeats show background levels of transcription (e.g. *Copia* and *Gypsy-7*), emphasizing that nascent transcription at the centromere occurs primarily at a subset of elements. Collectively, these analyses show that the *Drosophila* centromeres are actively transcribed and that *Jockey-3* in particular contributes significantly to the overall transcription occurring in these regions.

### *Jockey-3* transcripts localize to metaphase centromeres

*Jockey-3* is the only element that is transcribed at all five *Drosophila* centromeres (**Fig. 1D**). To examine the subcellular localization of *Jockey-3* transcripts in *D. melanogaster*, we designed strand-specific probes for single-molecule RNA Fluorescence *In Situ* Hybridization (smRNA FISH, henceforth RNA-FISH); one set detects sense transcripts targeting the 5 ’region of *Jockey-3*, spanning ORF1, and the other targets the 3 ’region, spanning the reverse-transcriptase domain within ORF2 (referred to as ORF1 and ORF2 probes; **Fig. 2A**). We also generated a reverse-complement set of the ORF2 probe to detect antisense transcripts (ORF2 anti). Each of the probe sets is made up of individual oligos that target both centromeric and non-centromeric *Jockey-3* (ORF1 = 44 oligos; ORF2 = 45 oligos). Several *Jockey-3* insertions across the genome are targeted by five or more probes, and are thus expected to produce RNA-FISH signal if sufficiently expressed, but centromere contigs are the regions targeted the most because 63% of *Jockey-3* copies are centromeric ((24); **Table 2 and Table S3**). Specifically, the ORF2 probe is expected to target primarily the *Jockey-3* copies on centromere X, Y, 3, and 4, while the ORF1 probe is expected to target those from centromere X, Y, 2, and 4.

**Figure 2.**
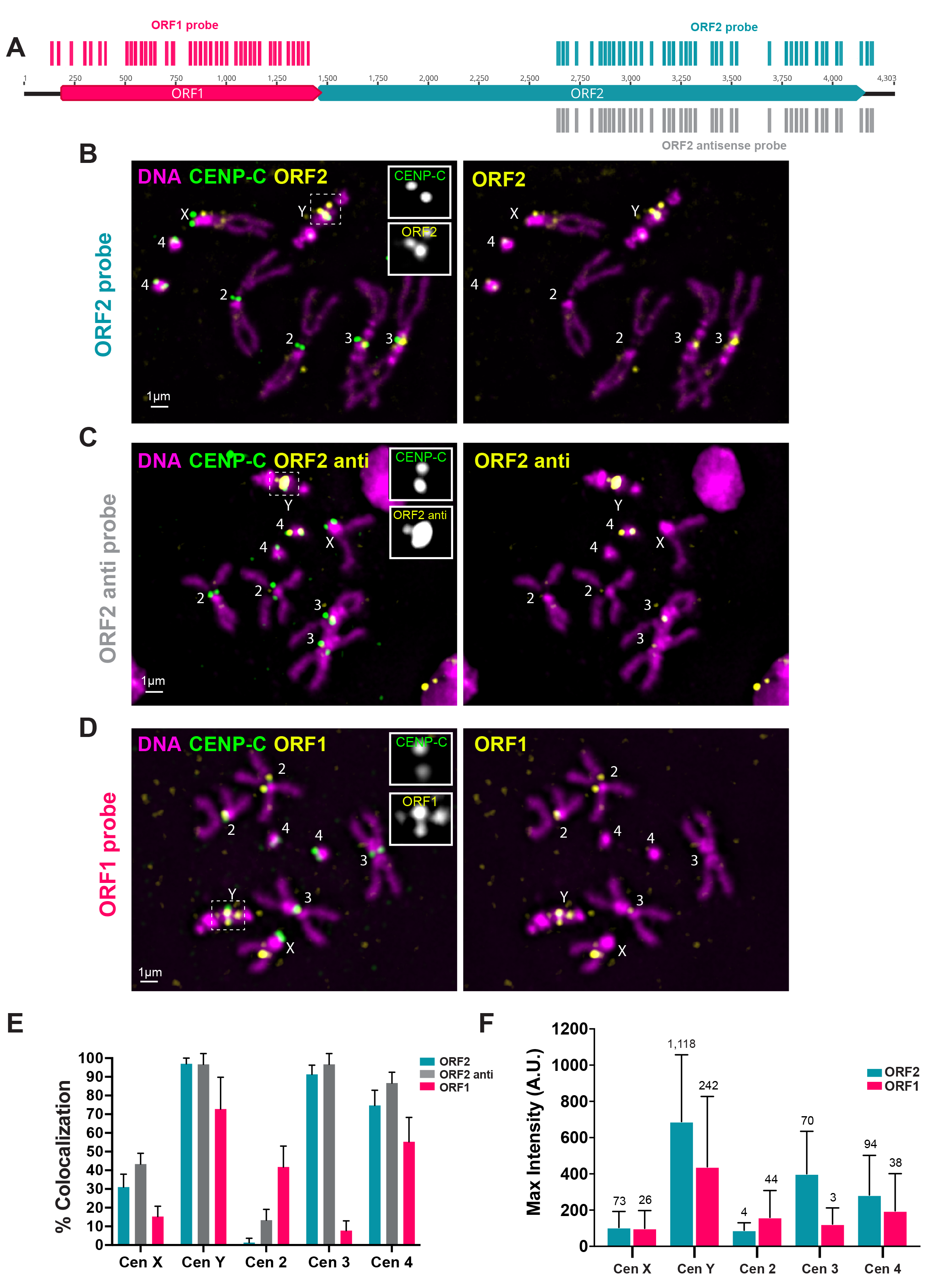
*Jockey-3* transcripts localize to metaphase chromosomes. **A** Diagram of *Jockey-3* showing base-pair position, predicted protein domains, and coverage of ORF1 (magents) and ORF2 (teal) probe sets. **B-D** Representative iso-1 male larval brain metaphase spreads. Chromosomes are stained with DAPI (magenta), RNA-FISH for *Jockey-3* ORF2 (B), ORF2 sense (C), and ORF1 (D) probes and IF for CENP-C (green). The images on the left show the merged channels and a grayscale 1.5x zoom inset for the Y centromere. The images on the right show DAPI and RNA FISH signals. **E** Graph for the percent of mitotic chromosomes showing colocalization between CENP-C and *Jockey-3* RNA FISH signal. ORF2 (N=3 brains, n=83 spreads), ORF2 anti (N=3 brains, n=28 spreads), and ORF1 (N=4 brains, n=69 spreads). **F** Maximum fluorescence intensity plot of centromeric *Jockey-3* RNA FISH signal. ORF2 probe (N= 1 brain, n=30 spreads) and ORF1 (N=1 brain, n=30 spreads). The numbers shown above each bar indicate the number of hits predicted to have complementarity with the corresponding probe set. A.U. stands for arbitrary units.

**Table 2.** Summary of RNA-FISH probe sequences centromere mapping. Table showing the total number of *Jockey-3* RNA-FISH probes predicted to bind to each centromeric and non-centromeric contigs. Full mapping across the genome is shown in Table S3.

| RNA-FISH probes mapping | ORF2 | ORF2 antisense | ORF1 |
| --- | --- | --- | --- |
| Cen X | 73 | 73 | 36 |
| Cen Y | 1117 | 1117 | 242 |
| Cen 2 | 4 | 4 | 44 |
| Cen 3 | 70 | 70 | 3 |
| Cen 4 | 130 | 130 | 38 |

We combined RNA-FISH for *Jockey-3* with immunofluorescence (IF; RNA-FISH/IF) for the centromere protein CENP-C which, unlike CENP-A, is retained on acid-fixed metaphase spreads from larval brain squashes. As a positive control for RNA-FISH, we used a smRNA-FISH probe targeting the *Rox1* non-coding RNA, which coats the X chromosome in males ((42); **Figure S4**). We observed transcripts labeled by the ORF2 probe co-localizing with CENP-C at the X, Y, 3rd, and 4th centromeres (**Fig. 2B, E and S5**), consistent with where these probes sequences map in the assembly (**Table 2 and Table S3**). We also observed co-localization of ORF2 antisense *Jockey-3* transcripts with CENP-C at the same centromeres (**Fig. 2C, E** and **Fig. S5**), indicating the simultaneous presence of both sense and antisense transcripts also shown by our transcript analyses (**Figs. 1 and S3**). Transcripts labeled by the ORF1 probe co-localized with centromeres X, Y, 2, and 4 (**Fig. 2D, E** and **S5**), again consistent with our predictions based on our mapping data (**Table 2** and **Table S3**).

The Y centromere is the only centromere containing full-length copies of *Jockey-3* and full-length copies show the highest levels of nascent transcription compared to other centromeres (**Fig. 1C**); thus it is not surprising that the Y displays co-localization between CENP-C and all three probe sets most consistently. In contrast, other chromosomes show more variability in signal detection (**Fig. 2E** and **S5**). In general, the frequency with which we observe co-localization between *Jockey-3* transcripts and CENP-C correlates with the number of probes targeting *Jockey-3* at each particular centromere, with centromere Y being targeted by the most probes overall due to this centromere containing 197/329 total *Jockey-3* copies in the genome ((24); **Figure 2E**, **Table 1-2** and **Table S3**). Maximum fluorescence intensity measurements for individual mitotic centromeres followed the same trend, with stronger signal detected on the Y (**Fig. 2F**). All five centromeres– including centromere 2, which contains only two *Jockey-3* fragments next to one another– show colocalization with at least one *Jockey-3* probe set. These findings confirm that truncated as well as full-length centromeric *Jockey-3* copies are active, consistent with our transcriptional profiles (**Figs. 1 and S3**). We also confirmed the localization of *Jockey-3* transcripts at metaphase centromeres in mitotic cells from ovaries and *Drosophila* Schneider cells (S2 cells; **Fig. S6 A-B**), confirming that this localization pattern is not unique to larval brain tissues. Furthermore, we performed RNA-FISH/IF on larval brains from *Drosophila simulans*, which diverged from *D. melanogaster* 2.5 million years ago (25) and whose centromeres are enriched in *Jockey-3* (24, 27). We observed centromeric foci for *Jockey-3* ORF2 at all mitotic centromeres, indicating that *Jockey-3* expression and transcripts localization is conserved in this species (**Fig. S6C**).

To ensure that the signal we observed with our *Jockey-3* probe sets corresponds to RNA and not DNA, we compared staining patterns between RNA and DNA-FISH protocols on brain squashes for the *Jockey-3* ORF2 probe and for a DNA-FISH OligoPaint targeting a 100-kb subtelomeric region of chromosome 3L band 61C7 (24). Using our RNA-FISH protocol, we could only detect the signal for *Jockey-3* produced by the ORF2 probe, while with our DNA-FISH protocol (which includes a DNA denaturation step and hybridization in the presence of an RNase cocktail) we only detected signal for the OligoPaint (**Fig. S7**). These experiments confirm that the *Jockey-3* signal shown in **Fig. 2B** corresponds to RNA and not DNA. Treatment with RNase H (which degrades DNA/RNA hybrids) post-hybridization dramatically reduced the signal intensity of *Jockey-3* foci, indicative of degraded DNA probe/RNA hybrids. We also observed a reduction in *Jockey-3* fluorescence when we performed a pre-incubation with an RNase cocktail expected to degrade single stranded RNA prior to RNA-FISH (**Fig. S8**). Together, these controls indicate that the *Jockey-3* transcripts we detect at centromeres with our RNA FISH protocol are *Jockey-3* single stranded transcripts.

In addition to localizing to centromeres, *Jockey-3* transcripts also localized to non-centromeric foci on all mitotic chromosomes with the exception of chromosome 4. On average, we observed 1 non-centromeric *Jockey-3* focus per mitotic spread, with a subset of cytological regions displaying foci more frequently than others (*e.g.* middle of XL; **Fig. S9**). Due to gaps in our genome assembly and the limited resolution that can be obtained by microscopy, it was not possible to determine to which *Jockey-3* copies these foci correspond.

Centromeric *Jockey-3* foci were also present in interphase cells from larval brains, ovaries, and S2 cells (**Fig. S10A-C**). On average, larval brains interphase cells displayed <1 *Jockey-3* focus co-localizing with CENP-C, versus 2-3 non-centromeric foci (**Fig. S10D**). Overall, mitotic cells display approximately 3 times more *Jockey-3* foci than interphase ones (**Fig. S10E**). Remarkably, only 15% of interphase cells display 2 or more *Jockey-3* foci co-localizing with CENP-C versus 93% of mitotic cells (**Fig. S10F**). *Drosophila* centromeres are often found clustered together in interphase, which might in part account for this difference. However, PRO-seq and RNA-seq data from larval brains, which reflect primarily the transcriptional state of interphase cells, show low coverage of Jockey-3 transcripts at the centromere islands (**Fig. S3**), consistent with overall lower transcription occurring at the centromere in interphase compared to mitosis. We note that the non-centromeric *Jockey-3* foci observed in interphase could reflect transcripts that remain associated in *cis* or unbound nuclear RNAs.

Lastly, to expand on our RNA localization studies, we designed smRNA-FISH probes for another centromeric non-LTR element, *Doc*, which is found within centromere X and 4 and that shows expression (**Fig. 1**). We performed smRNA-FISH/IF on mitotic and interphase cells from larval brains squashes. Unlike *Jockey-3*, *Doc* transcripts were not detectable at the centromeres in metaphase, although the signal was visible in a few interphase cells, where it co-localized with one CENP-C focus (**Fig. S11**). We conclude that not all centromeric retroelements produce transcripts that localize to centromeres in metaphase.

### *Jockey-3* transcripts co-localize with their cognate sequences in *cis*

Studies in human and *Drosophila* cultured cells and in *Xenopus* egg extracts reported that different centromere and pericentromere-derived repeat transcripts can localize to centromeres either in *cis* (*i.e.* at the locus of origin; (9, 21)) or in *trans* (*i.e.* to all centromeres whether or not they contain complementary sequences (12, 23)). Two observations from our data so far point towards *cis* localization of *Jockey-3* transcripts at the centromere. First, the centromeric signal intensity for *Jockey-3* RNA-FISH is positively correlated with the number of probes targeting that centromere (**Fig. 1F** and **Table 1** and **S1**), whereas with *trans* localization, a more uniform signal intensity would be expected, irrespectively of the DNA composition of each centromere. Second, *Drosophila* centromere 2 contains two fragments of *Jockey-3*, one targeted by only 4 out of 44 probes in the ORF2 set and the other targeted by 44 out of 45 probes in the ORF1 set (**Fig. 3A** and **Table 2**) and we observe robust RNA-FISH signal nearly exclusively with the one targeting ORF1 (**Fig. 3B** and **1E**). Conversely, centromere 3 *Jockey-3* copies are targeted primarily by ORF2 probes and indeed we observe strong centromeric signals for ORF2 but not ORF1. These observations indicate that RNAs emanating from *Jockey-3* copies colocalize with their cognate DNA sequences in *cis*.

**Figure 3.**
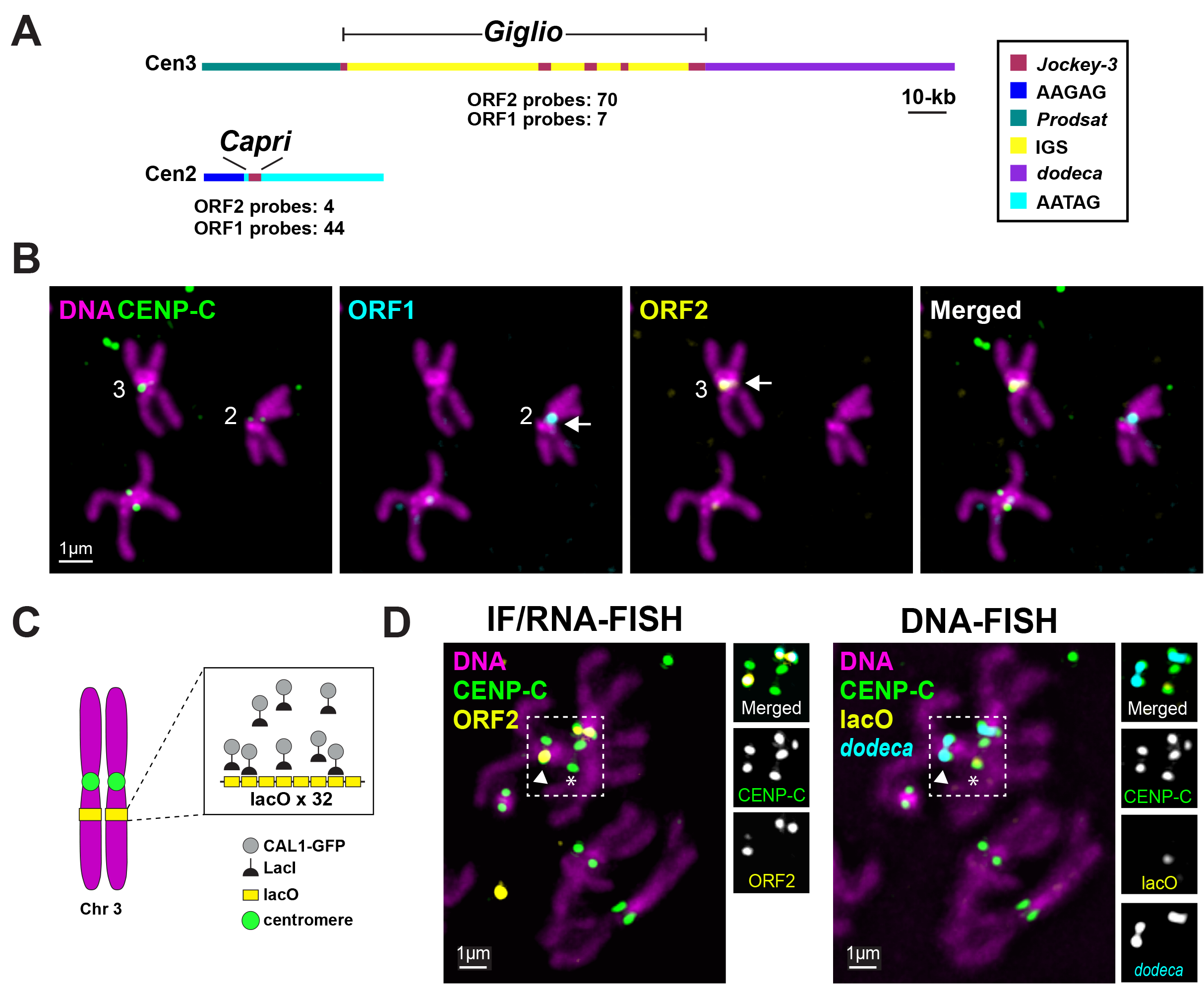
*Jockey-3* transcripts co-localize with their cognate sequences in *cis*. **A** Schematic showing the organization of centromere 3 (top) and 2 (bottom) and the number of probes from the ORF1 and the ORF2 (both sense) predicted to bind to the *Jockey-3* elements therein. **B** Representative spread from RNA-FISH/IF in iso-1 flies showing the presence of *Jockey-3* signal for the ORF2 (yellow) at the centromere of chromosome 3 (arrowhead) and for the ORF1 (cyan) at the centromere of chromosome 2. CENP-C (green) and DNA stained with DAPI (magenta). Bar 1µm. **C** Schematic showing the *de novo* centromere system for chromosome 3 (lacO 3^peri^). Progeny containing one lacO chromosome 3, UAS-CAL1-GFP-LacI, and elav-GAL4 were analyzed by sequential IF/RNA/DNA FISH. **D** Sequential IF/RNA (left)/DNA-FISH (right) on larval brain metaphase spreads of *de novo* centromere progeny (CAL1-GFP-LacI; lacO 3^peri^) showing *Jockey-3* transcripts (3 ’probe; yellow) overlapping with the endogenous centromere 3 (yellow arrowhead) but not the *de novo* centromere on lacO (asterisk). CENP-C is a centromere marker (green), *dodeca* is a satellite specific for centromere 3 (cyan). The lacO array DNA FISH is shown in yellow in the right panel. Bar 1µm. N=6 brains (3 males, 3 females), n=90 cells total.

To more robustly test if *Jockey-3* transcripts can localize in *trans* to other centromeres, we asked if *Jockey-3* transcripts can be detected on a *de novo* centromere formed on DNA devoid of any centromere-associated repeats. We used a previously developed LacI/lacO system that efficiently forms ectopic centromeres *in vivo* via the tethering of the CENP-A assembly factor CAL1, fused to GFP-LacI, to a 10-kb lacO array inserted at the pericentromere of chromosome 3 (43). We analyzed a total of 89 metaphase spreads from 3 male larval brains by IF/RNA-FISH with anti-CENP-C antibodies and the ORF2 probe and, after imaging, performed sequential DNA-FISH to confirm the location of lacO in the same spreads. We found that, while robust localization of *Jockey-3* ORF2 transcripts at endogenous centromere 3 was clearly visible, *Jockey-3* signal was nearly never observed at the ectopic centromere on lacO (1/90 showed weak signal on one sister; **Fig. 3B-C**). Together, these findings are consistent with *Jockey-3* transcripts remaining associated with the DNA sequences they originated from, similarly to what was reported for centromeric alpha-satellite transcripts in human cells (9).

### Knockdown of *Jockey-3* RNA does not negatively affect normal centromere function

Knock-downs of alpha-satellite transcripts (44) and of a LINE-1 element associated with a neocentromere (17) lead to a decrease in the levels of CENP-A from the (neo)centromeres these transcripts originate from, suggesting they play a localized role in centromere maintenance or stability. In contrast, in *S. pombe,* centromere-derived transcripts are rapidly degraded by the exosome and are thus unlikely to play such a structural role, but rather appear to be byproducts of centromere transcription (29).

To test the possibility that *Jockey-3* transcripts themselves play a role in centromere integrity, we designed a short-hairpin (sh) to target *Jockey-3* RNA for degradation via *in vivo* RNA interference (RNAi). As *Jockey-3* copies are heavily polymorphic in sequence and length, no single sh can target the majority centromeric or genomic copies. We therefore designed a sh targeting the RT domain in ORF2, which is present in ∼27% of *Jockey-3* insertions in the genome, targeting as many centromeric and non-centromeric copies as possible (**Fig. 4A**), and generated transgenic flies expressing the sh-*Jockey-3* under a GAL4 UAS promoter.

**Figure 4.**
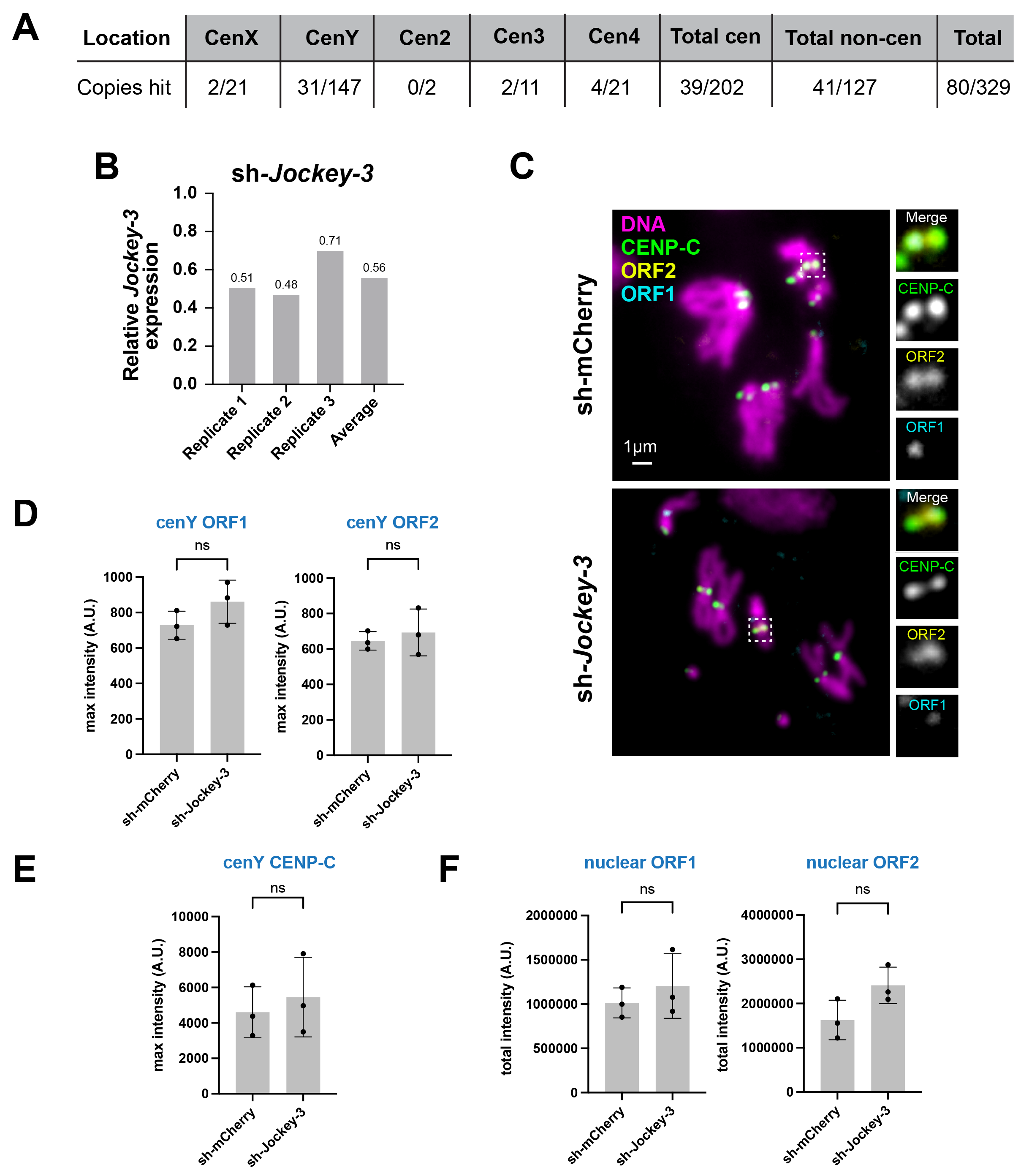
Knockdown of *Jockey-3* RNA does not negatively affect normal centromere function. **A** Table showing centromeric and non-centromeric *Jockey-3* copies targeted by the sh-*Jockey-3* over the total number of *Jockey-3* copies. Targets with up to 3 mismatches are included. **B** Efficiency of *Jockey-3* knockdown determined by RT-qPCR normalized to Rp49 and set relative to sh-mcherry control in elav-GAL4 male larval brains. The average of three biological replicates are shown. The primers used here capture 72/329 (ORF2 RT primer set) *Jockey-3* copies throughout the genome and 72/80 targeted by the sh, 32 of which are centromeric copies (two on X, 27 on the Y, 2 on the 3rd, and 3 on 4th chromosome). **C** Representative images of mitotic spreads from larval brains expressing sh-mcherry control and sh-*Jockey-3* stained by IF/RNA-FISH with CENP-C antibodies (green) and *Jockey-3* ORF1 (cyan) and ORF2 (yellow) probes. Insets show a zoomed image of the centromeres in the box. Bar 1µm. **D** Quantification of *Jockey-3* ORF2 and ORF2 RNA-FISH signals at the Y centromere. Bar graphs show the average fluorescence intensity for *Jockey-3* ORF2 and ORF1 at the Y centromere from sh-mcherry and sh-*Jockey-3* (unpaired t-test, p>0.05 for both the *Jockey-3* ORF2 and ORF2, N=3 brains, n=25 Y centromeres/brain). A.U. stands for arbitrary units. **E** Quantification of CENP-C signals at the Y centromere. The bar graph shows the average fluorescence intensity for CENP-C at the Y centromere from sh-mcherry and sh-*Jockey-3* (unpaired t-test, p>0.05, N=3 brains, n=25 Y centromeres/brain). A.U. stands for arbitrary units. **F** Quantification of *Jockey-3* ORF2 and ORF2 RNA-FISH signals in the total interphase cell nucleus. Bar graphs show the average fluorescence intensity for *Jockey-3* ORF2 and ORF1 in the cell nucleus from sh-mcherry and sh-*Jockey-3*. (unpaired t-test, p>0.05 for both *Jockey-3* ORF2 and ORF2, N=3 brains, n=25 Y centromeres/brain). A.U. stands for arbitrary units.

To verify the effectiveness of the knock-down, we induced sh-*Jockey-3* expression under the neural elav-GAL4 driver, isolated total RNA from larval brains, and measured *Jockey-3* expression by RT-qPCR, using primers mapping outside of the sh-*Jockey-3* target. These primers capture 72/80 *Jockey-3* copies targeted by the short hairpin, including 2 centromeric copies on the X, 27 on the Y, 2 on the 3rd, and 3 on 4th chromosome, all of which were confirmed as expressed by PRO-seq. Across three biological replicates, we found that sh-*Jockey-3* expression was reduced by ∼44% in sh-*Jockey-3* compared to a sh-mcherry control (**Fig. 4B**). However, measurements of the RNA-FISH signal intensity showed no significant change for *Jockey-3* ORF1 or ORF2 at the Y centromere in metaphase (**Fig. 4C-D**). Similarly, we did not observe a decrease in CENP-C intensity at the Y centromere (**Fig. 4E**), which would have been indicative of a centromere assembly defect, nor did we detect an increase in aneuploidy (N=3 brains, n=25 spreads each, 1.33% aneuploid in sh-*Jockey-3* versus 6.7% in control, p=0.2).

RNAi based knockdowns typically affect transcripts post-transcriptionally and their effectiveness in knocking down nuclear RNAs is unclear (discussed in (45)). To determine if the nuclear pool of *Jockey-3* transcripts is reduced upon RNAi, we quantified the total nuclear fluorescence intensity of *Jockey-3* in interphase larval brain cells and found no significant change compared to the control (**Fig. 4F**), suggesting that the decrease in expression observed by RT-qPCR (**Fig. 4B**) reflected changes in the cytoplasmic pool of *Jockey-3*. An alternative explanation is that the *Jockey-3* copies not targeted by the knockdown supply sufficient nuclear RNA signal to obfuscate any reductions caused by the depletion. Nonetheless, consistent with the lack of mitotic defects, expression of the hairpin under the eyeless-GAL4 driver in adult eyes did not cause any disruptions to eye morphology compared to the control (data not shown). We also observed similar progeny viability and fertility in flies expressing sh-*Jockey-3* compared to controls (data not shown). These findings suggest that the cytoplasmic pool of *Jockey-3* RNA is not important for centromere integrity, chromosome segregation, or viability. However, given that this approach does not target all expressed *Jockey-3* copies, we cannot rule out that nascent *Jockey-3* RNA may play a role as a *cis*-acting non-coding RNA at centromeres.

### CENP-A chromatin profiling reveals a link between *Jockey-3* transcription and CENP-A association

*Jockey-3* is the most enriched repeat in CENP-A chromatin immunoprecipitations (24) and is present at both centromeric and non-centromeric regions of the genome (**Table S1**). However, the non-centromeric occupancy of *Drosophila* CENP-A and its relationship with non-centromeric *Jockey-3* copies has not been explored. Furthermore, we do not know if the presence of CENP-A and the transcriptional activity of *Jockey-3* are correlated. To investigate these questions, we identified significant CENP-A peaks using CUT&Tag (46) from 0-12h embryos, and mapping the resulting sequencing data to the heterochromatin-enriched genome assembly (24). We identified the expected five centromeric CENP-A domains (**Fig. S12**; (24)) along with 333 non-centromeric domains (**Table 3**; **Table S4**). These non-centromeric CENP-A domains were smaller on average and contained lower CENP-A signal intensity than the centromeric ones (**Fig. 5A-B**). Lower CENP-A signal of ectopic compared to centromeric CENP-A was also previously reported for human HeLa cells (47). Next, we examined whether the transcription of *Jockey-3* copies correlated with CENP-A occupancy. There are 202 copies of *Jockey-3* that fall within a centromeric CENP-A domain, 26 that fall within a non-centromeric CENP-A domain, and 101 that fall in neither (**Table S6**). We found that, while 36% of *Jockey-3* copies within the centromeric CENP-A domains are expressed, this percentage increases to 96% for *Jockey-3* copies at non-centromeric CENP-A regions. Expression of *Jockey-3* copies not CENP-A associated is also high at around 60% (**Fig. 5C; Table S1** and **Table S5**). When we compared all CENP-A associated *Jockey-3* copies with all non-CENP-A associated ones, the difference in the percentage of active *Jockey-3* elements is only 43% versus 62%, respectively (**Fig. 5D; Table S1** and **Table S5**). We conclude that although there is an enrichment of *Jockey-3* elements associated with CENP-A versus not (228/329, or 69%; **Fig. 5E** and **Table S6**), the expression of *Jockey-3* in embryos appears to be independent of its association with CENP-A. However, when we consider only full-length *Jockey-3* copies, which are the most highly expressed copies in the genome (**Fig. 1B**), we see a strong and positive correlation between the association with CENP-A and active transcription (**Fig. 5F; Table S1**), regardless of centromeric location. After breaking down the data by where all full-length *Jockey-3* copies are located (Y centromere, non-centromeric regions, or non-CENP-A associated regions), it is clear that CENP-A association, irrespective of centromeric location, is correlated with higher transcription (**Fig. 5G**). From these analyses, we conclude that full-length *Jockey-3* copies are more highly expressed when coupled with CENP-A chromatin.

**Figure 5.**
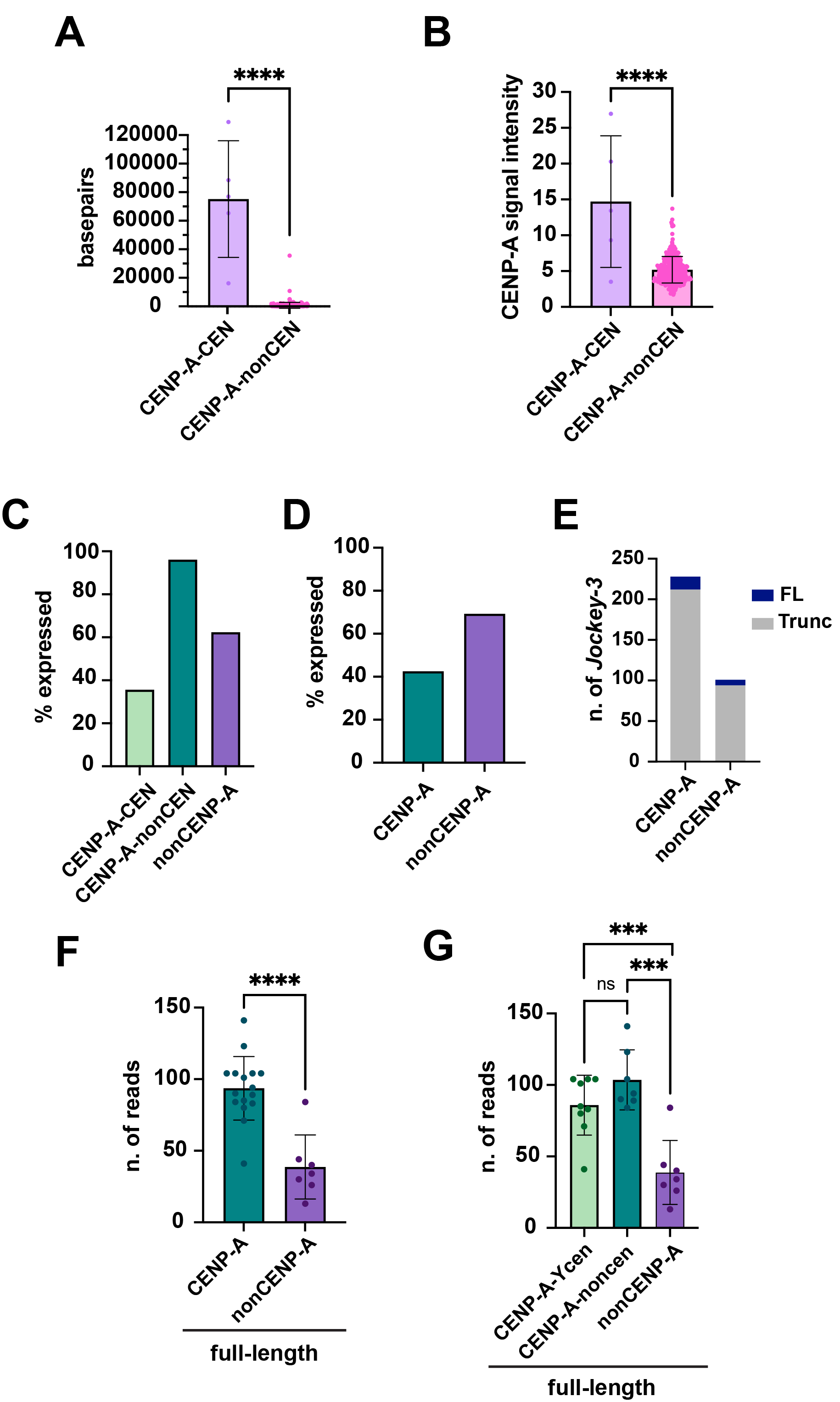
Relationship between CENP-A occupancy and transcription at centromeric and non-centromeric *Jockey-3* insertions. **A** Scatter boxplot showing CENP-A domain size (in base pairs) between centromeric (n=5) and non-centromeric (n=333) loci based on MACS2 peak calls from CUT&Tag data. Statistical significance was determined with unpaired t-test (****; p < 0.0001; Student’s t-test). Error bars show the standard deviation. **B** Scatter boxplot showing CENP-A peak signal intensity between centromeric and non-centromeric loci based on MACS2 peak calls from CUT&Tag data. Signal intensity was averaged across each CENP-A domain. Statistical significance was determined with unpaired t-test (****; p < 0.0001; Student’s t-test). Error bars show the standard deviation. **C** Bar graph illustrating the proportion of *Jockey-3* copies expressed per group, where groups are based on CENP-A and centromeric association. PRO-seq mapping was done with Bowtie 2 default “best match” using paired-end reads, post-deduplication. Expression is defined as having at least two PRO-seq read overlaps. **D** Same as shown in **C**, except *Jockey-3* copies found within CENP-A domains (regardless of centromeric association) are combined into one group (“CENP-A”). **E** Distribution of *Jockey-3* copies as a stacked bar graph. Copies are grouped by whether they are found within CENP-A domains (regardless of centromeric association) or outside CENP-A domains, as well as their status as a full-length (blue) or truncated elements (gray). **F** PRO-seq read density scatter boxplot of full-length *Jockey-3* copies comparing those found within CENP-A domains (centromeric and non-centromeric) and those found outside CENP-A domains. Mapping was done with Bowtie 2 default “best match” using paired-end reads, post-deduplication. Statistical significance was determined with unpaired t-test (****; p < 0.0001). Error bars show the standard deviation. **G** Same as shown in **F**, except full-length *Jockey-3* copies found within CENP-A domains are split by centromeric (present only within the Y centromere) or non-centromeric locations. Unpaired t-tests (Student’s t-test) were performed between each group (***, p < 0.001; ns (non-significant), p > 0.05). Error bars show the standard deviation.

**Table 3.** Summary of the CENP-A domains and associated *Jockey-3* insertions. Table showing the distribution of CENP-A domains classified as centromeric vs. non-centromeric and the proportion that contains copies of *Jockey-3*.

| CEN designation | # CENPA domains | # CENPA domains containing *Jockey-3* |
| --- | --- | --- |
| CEN | 5 (1/chromosome) | 5 (100%) |
| nonCEN | 333 | 43 (12.91%) |

It is noteworthy to point out that both PRO-seq and CUT&Tag were performed on nuclei from embryos and thus reflect the transcriptional and chromatin profiles of primarily interphase cells. In contrast, the observation that the detection of *Jockey-3* RNA-FISH signal is more frequent at centromeres compared to non-centromeric locations came from metaphase chromosomes (**Fig. S10C**). Even though we cannot directly test this by PRO-seq on mitotic cells, we infer that the proportion of *Jockey-3* transcripts emanating from centromeres versus non-centromeric regions is likely to be higher in mitosis.

### Recent *Jockey-3* insertions are found more frequently within CENP-A chromatin and are more expressed

In *Drosophila*, *Jockey-3* shows weak insertional bias for the centromere (41), but whether such preference relies on specific centromeric sequence features or on the presence of CENP-A is unknown. The observation that *Jockey-3* is also enriched at the centromeres of *D. simulans* (24, 27), even though this species contains widely divergent centromeric repeats (26, 27), suggests that such insertion bias is unlikely to be mediated by DNA sequence preference. If *Jockey-3* preferentially transposes within centromeres through recognition of CENP-A chromatin, we would expect recent insertions to be enriched within both centromeric and non-centromeric CENP-A domains. To test this possibility, we calculated the percentage of young *Jockey-3* insertions (<1% divergence from *Jockey-3* consensus; **Table 1**; (41)) that overlap with CENP-A domains and compared it to the percentage found at non-CENP-A containing regions of the genome. Interestingly, we found that 80% of young copies (34/42) are found in genomic regions that overlap with CENP-A domains, compared to 20% in non-CENP-A containing regions (**Fig. 6A**). Considering that CENP-A domains make up a small percent of the genome, this is a dramatic enrichment. Of these 34 CENP-A-associated *Jockey-3* copies, 13 are centromeric and 21 non-centromeric, consistent with the hypothesis that the retroelement targets CENP-A chromatin for reinsertion irrespective of its centromeric or non-centromeric location. In contrast, old *Jockey-3* insertions (>1% divergence from *Jockey-3* consensus; **Table 1**; (41)) are disproportionately associated with centromeric CENP-A domains rather than non-centromeric ones. One possible explanation for this observation is that non-centromeric CENP-A domains are more dynamic over evolutionary time than centromeres and thus, as retroelement insertions in those regions age, they end up no longer being CENP-A associated.

**Figure 6.**
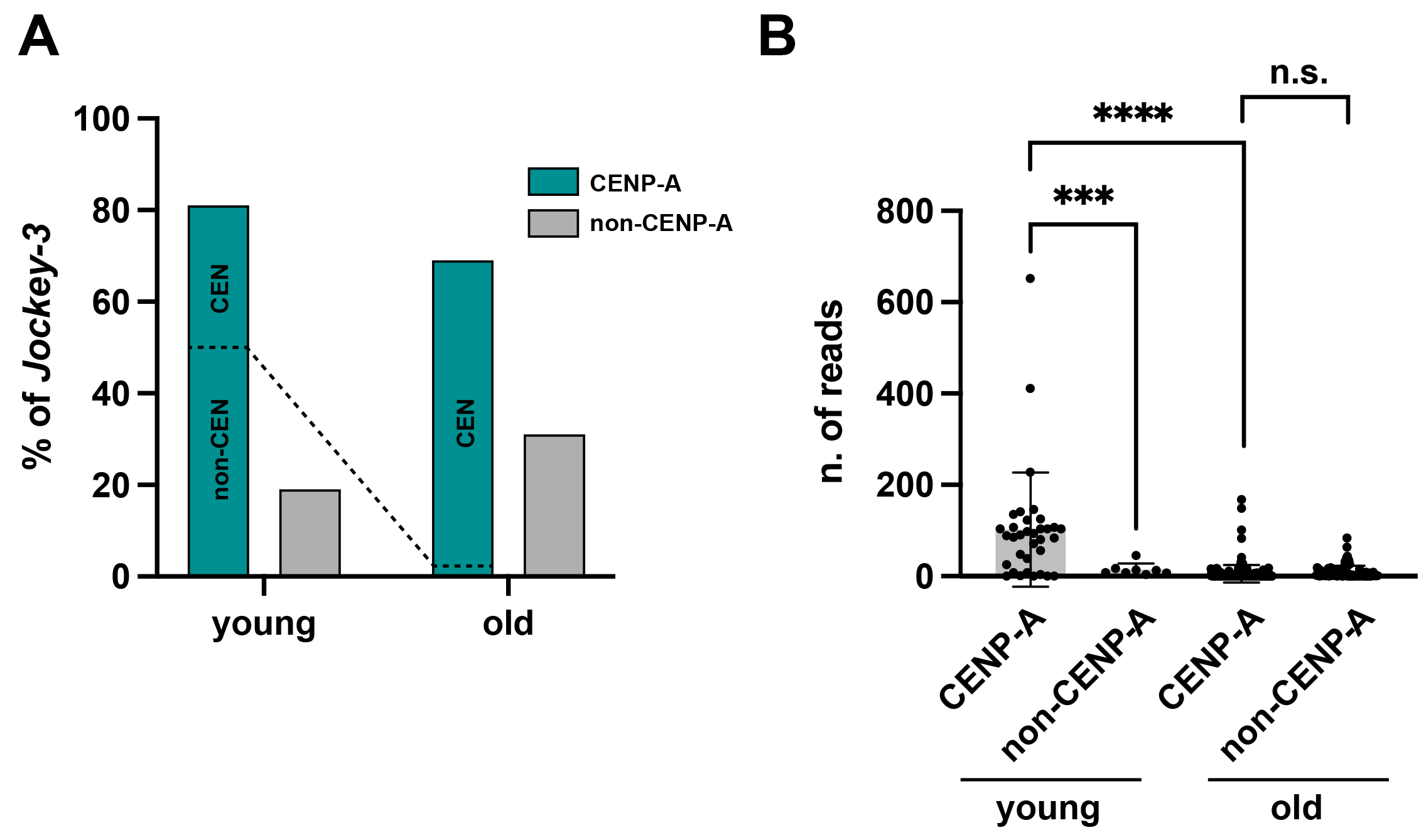
Recent *Jockey-3* insertions are found more frequently within CENP-A chromatin and are more expressed. **A** Percentage of young *Jockey-3* copies (<1% divergence from consensus) found within CENP-A domains, designated as centromeric (CEN) and non-centromeric (non-CEN), versus non-CENP-A regions identified by CUT&Tag. 61% of young insertions (21/34) are at non-centromeric CENP-A domains (non-CEN) compared to 38% (13/34) centromeric (CEN). **B** PRO-seq read counts mapping to young versus old Jockey-3 copies with Bowtie 2 default “best match”, post-deduplication (****, p<0.0001, unpaired t-test).

The presence of CENP-A on full-length *Jockey-3* copies is associated with higher transcription (**Fig. 5G**). We hypothesized that *Jockey-3* preferentially inserts within CENP-A chromatin to increase its chance of being expressed. If this were the case, we would expect recent insertions to be more highly expressed if associated with CENP-A than not. We counted the number of PRO-seq reads mapping to CENP-A associated and non-CENP-A associated *Jockey-3* insertions classified as young or old (**Table 1**) and found that newer insertions within CENP-A chromatin are significantly more expressed than those at non-CENP-A domains (**Fig. 6B**). Older insertions are overall less expressed than young ones. Interestingly, young CENP-A associated copies, which are primarily non-centromeric (**Fig. 6A**) are also more expressed than their older counterparts, which are primarily centromeric. However, the centromeric *Jockey-3* copies are also largely truncated, which we showed are generally less transcribed (**Fig. 1B**). Collectively, these observations suggest a model where *Jockey-3* has evolved the ability to target CENP-A for insertion to promote its expression. Due to its role at centromeres and its requirement to be transcriptionally permissive, CENP-A chromatin may be spared by genome-defense mechanisms that target transposons for silencing, providing a protective environment for *Jockey-3*.

### *lacO* transcription is coupled with *de novo* centromere formation

All our data so far points to a correlation between CENP-A chromatin and *Jockey-3* expression. Therefore, we next investigated if DNA associated with *de novo* centromeres, which lack *Jockey-3* or other centromere repeats, is also transcribed. In *Drosophila* S2 cells and flies *de novo* centromeres are efficiently formed when the CENP-A chaperone CAL1 is fused to GFP-LacI and tethered to a lacO array inserted within the genome (43, 48). Upon its tethering to the lacO array in S2 cells, CAL1, alongside the elongation factor FACT and RNA polymerase II, initiate transcription of non-endogenous sequences belonging to the inserted lacO array (16).

To determine if the DNA associated with a *de novo* centromere becomes transcribed *in vivo*, we used an oligo lacO probe to detect lacO-derived transcripts by RNA-FISH in larval progeny expressing CAL1-GFP-LacI or a GFP-LacI control under the neural elav-GAL4 promoter and heterozygote for a pericentric 10-kb lacO array inserted at 3L (3^peri^ at cytoband 80C4; (43)). Consistent with previous studies, expression of CAL1-GFP-LacI results in ectopic centromere formation at the 3^peri^ lacO array in more than 80% of spreads (43). We performed sequential IF-RNA/DNA-FISH on mitotic spreads from larval brains in elav-GAL4, CAL1-GFP-LacI and GFP-LacI/lacO expressing progeny. IF for CENP-C was used to identify active centromeres and lacO RNA-FISH allowed us to establish if transcripts are visible at ectopic centromeres. After imaging metaphase spreads, we processed the slides for DNA-FISH with the same lacO probe to identify the position of the lacO array. We also included a probe for the peri/centromeric satellite *dodeca* to identify the endogenous centromere 3s, and re-imaged the same mitotic spreads. We found that both GFP-LacI control spreads and CAL1-GFP-LacI/3^peri^ spreads display lacO RNA-FISH signal, but the latter show significantly higher frequency compared (**Fig. 7A-B**). In interphase, we found that there is no significant difference in lacO transcription frequency between CAL1-GFP-LacI/3^peri^ and GFP-LacI/3^peri^ in interphase cells (**Fig. 7C**), suggesting that the higher transcription frequency observed in CAL1-GFP-LacI/3^peri^ is specific to metaphase. To determine if lacO expression levels are different between GFP-LacI and CAL1-GFP-LacI/3^peri^ mitotic spreads, we measured lacO RNA fluorescence intensity for both genotypes and found that CAL1-GFP-LacI/3^peri^ displays higher lacO RNA signal intensity than the GFP-LacI/3^peri^ control (**Fig. 7D**). Collectively, these experiments demonstrate that although lacO is transcribed in the absence of an ectopic centromere, transcription is observed at a higher frequency and at higher levels when an ectopic centromere is present, suggesting that the formation of a *de novo* centromere stimulates local transcription. These results are consistent with previous reports in human neocentromeres (17, 19, 49) and *de novo* centromeres in S2 cells (16) showing increased transcription upon CENP-A chromatin formation at non-centromeric sites. They also further underscore the correlation between CENP-A deposition in mitosis and an increase in transcription.

**Figure 7.**
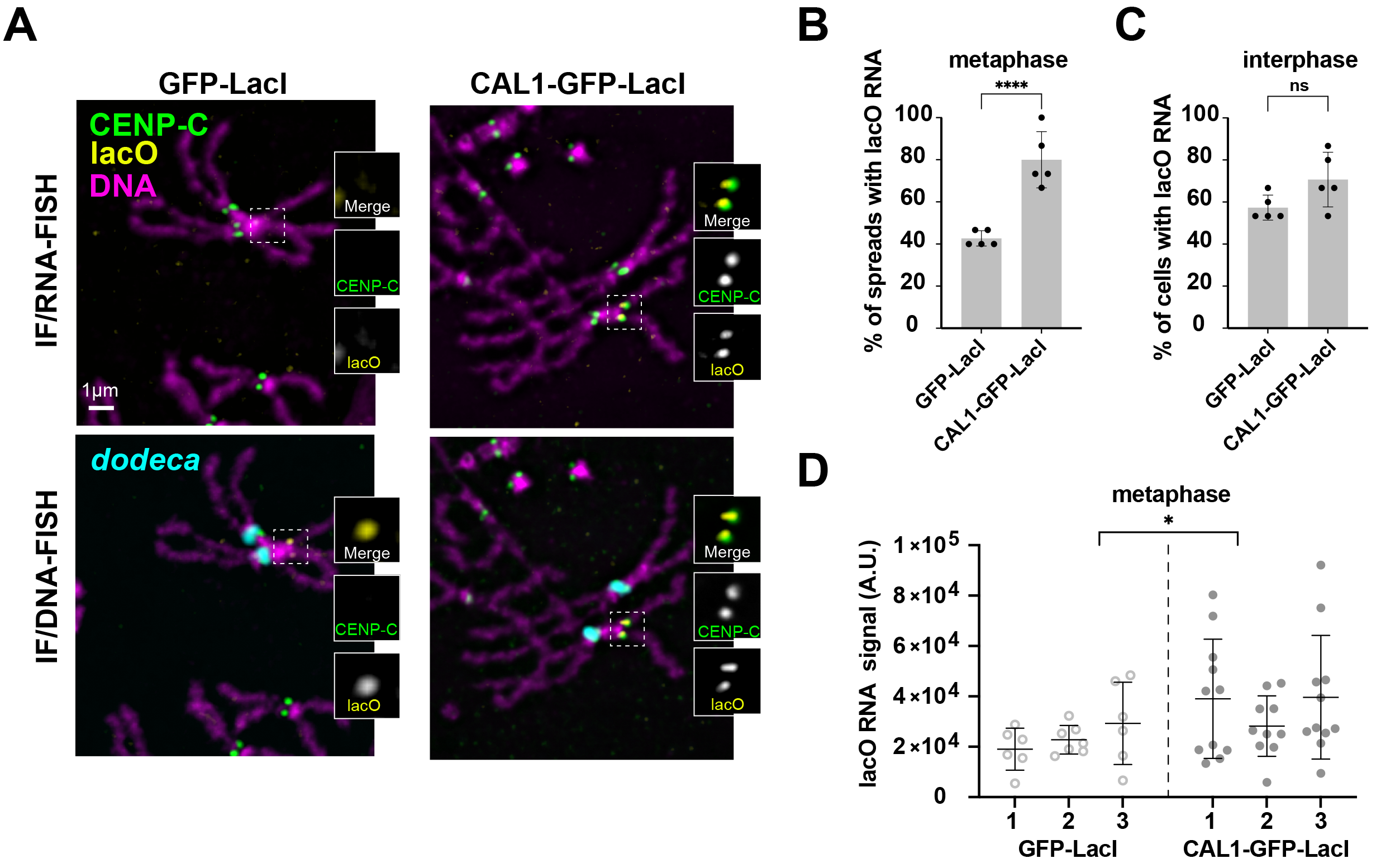
*lacO* transcription is coupled with *de novo* centromere formation. **A** Sequential IF/RNA/DNA FISH on larval brains from GFP-LacI and CAL1-GFP-LacI/3^peri^, both (lacO array at 3^peri^). IF for CENP-C is shown in green. RNA and DNA FISH with a lacO LNA probe are shown in yellow. DNA FISH for *dodeca* is shown in cyan. **B** Bar graphs showing the frequency of lacO transcription in GFP-LacI and CAL1-GFP-LacI/3^peri^ in metaphase and **C** in interphase (Fisher’s exact test, N = 5 brains, n = 15 spreads/brain). **D** Scatter plot showing the fluorescence intensity of lacO RNA-FISH in GFP-LacI and CAL1-GFP-LacI/3^peri^ in metaphase (nested t-test, N = 3 brains, n = 6-11 spreads/brain). A.U. stands for arbitrary units.

## Discussion

In this study, we examined the transcriptional landscape of *Drosophila* centromeres and identified the centromere-enriched retroelement *Jockey-3* as a key transcribed component across these regions. We found that *Jockey-3*, produces transcripts that accumulate at all mitotic centromeres, a localization that is conserved in *D. simulans*. In metaphase, *Jockey-3* transcripts remain associated with their cognate DNA sequences and do not diffuse to other native nor *de novo* centromeres. Metaphase is the cell cycle stage that coincides or precedes (depending on cell types and species) metazoan CENP-A deposition (50–56). A boost in transcription before or around the time of CENP-A deposition could prime chromatin by removing place-holder histone H3.3 (57) to allow the assembly of CENP-A nucleosomes. Consistent with this model, active RNA polymerase II (RNAPII) and/or transcriptional activity has been reported at metaphase centromeres in both *Drosophila* (21, 23) and human cell lines (9, 11, 30). In human cells, RNAPII is lost from chromosome arms upon cohesin degradation in prophase, yet persists at centromeres in metaphase where cohesin remains enriched until anaphase (58).

To inform on whether the act of transcription is important for CENP-A maintenance, previous studies used transient treatments with RNA polymerase inhibitors. In *Drosophila* S2 cells, transcriptional blockage destabilized the chromatin association of new CENP-A at centromeres (21). Somewhat surprisingly, RNA polymerase inhibitors injected into early *Drosophila* embryos did not result in a decrease in centromeric GFP-CENP-A signal intensity, which would be expected if transcription was required for *de novo* GFP-CENP-A deposition (59). However, it is unclear if CENP-A deposition during the rapid divisions occurring at this developmental stage involves eviction of place-holder histone H3.3.

There are 329 copies of *Jockey-3* in the *Drosophila* genome, 202 of which (61%) are found within the five centromere contigs (24, 41). Analyses of nascent transcripts reveal that the *Jockey-3* copies present within the centromeres are not expressed at higher levels than those found elsewhere in the genome –in fact, at least in interphase, *Jockey-3* elements within the centromeres are overall expressed at lower levels– suggesting that the expression of *Jockey-3* elements is not linked to their centromeric location. These results are consistent with studies in human RPE cells that showed that alpha-satellite transcripts are produced from both centromeric arrays and from arrays outside of the active human centromere region (9). It is possible that the accumulation of *Jockey-3* and other expressed repeats at the centromere might underscore selection for transcriptionally active elements in these regions to facilitate CENP-A chromatin maintenance.

Full-length *Jockey-3* copies contribute the most to overall *Jockey-3* transcription, and the majority of these full-length copies are found at non-centromeric loci (14/23). Interestingly, we find that the expression of these full-length *Jockey-3* copies is strongly positively correlated with CENP-A occupancy. Our PRO-seq profiles reflect nascent transcription in interphase, and at this cell cycle stage the co-localization of *Jockey-3* RNA signal with centromeres is detected less frequently and at fewer centromeres than in metaphase. In contrast, RNA-FISH on metaphase chromosomes reveals bright *Jockey-3* RNA foci primarily at centromeres. The observation that transcripts from the expressed centromere-associated retroelement *Doc* do not localize to metaphase centromeres, unlike those from *Jockey-3*, suggests that *Jockey-3* may have a unique ability for enhanced transcription during this stage. The RNA signal is especially strong on the mitotic Y centromere, which contains an abundance of expressed *Jockey-3* copies. It is interesting to note that the Y centromere also displays stronger CENP-A signal in spermatocytes and early embryos (60), consistent with the possibility that high levels of CENP-A may be linked to abundant *Jockey-3* expression and/or the retention of its RNA products.

While centromere-associated *Jockey-3* transcripts are visible with high frequency in metaphase, non-centromeric foci are more rare and certainly fewer than the 127 known non-centromeric *Jockey-3* insertions or the 14 full-length non-centromeric copies. In interphase too, the number of non-centromeric foci is much smaller than the number of non-centromeric *Jockey-3* copies. It is possible that different insertions alternate between active and inactive states. Alternatively, only a subset of full-length *Jockey-3* copies produce sufficient nascent transcripts to be detectable by RNA-FISH.

Our finding that *de novo* centromeres are coupled with transcriptional activation of the underlying DNA specifically in metaphase reinforces the model that CENP-A deposition and transcription go hand in hand. Our experiments do not distinguish between transcriptional activation of lacO being caused by CAL1 tethering, given that CAL1 is known to interact with RNAPII and FACT (16), or being linked to active CENP-A deposition. However, the latter possibility would be consistent with recent studies in human neocentromeres showing that neocentromere formation is associated with transcriptional activation and increased chromatin accessibility (18, 19).

The *Jockey-3* retroelement is enriched at the centromere compared to the rest of the genome in *D. melanogaster* and *D. simulans* (24, 27). How this retroelement has accumulated at centromeres over time remains a matter of speculation, but population studies show that low frequency polymorphic insertions, indicative of recent transpositional events, show a weak bias towards centromeres (61). Using divergence from the consensus to estimate the age of the element (61), we found that much of the most recent transposition events have occurred within regions containing CENP-A. Given that the majority of non-centromeric CENP-A domains do not overlap with a *Jockey-3* element, we speculate that it is *Jockey-3* that follows CENP-A rather than the other way around. Regardless of whether CENP-A or *Jockey-3* come first, recent *Jockey-3* copies are more transcribed than old ones, suggesting that a new insertion has the potential to affect CENP-A chromatin, which could result in its stabilization or its disruption.

The centromeres of three species within the *Drosophila simulans* clade–*D. simulans*, *D. mauritiana*, and *D. sechellia–*and *D. melanogaster*, display a remarkable turnover in sequence composition, suggesting the existence of a genetic conflict between satellites and retroelements (27). To ensure their own propagation through generations, these selfish genetic elements appear to compete for dominance at the centromere, a region with low recombination that can tolerate variation in sequence composition without loss of functionality. Since *Jockey-3* is targeted by piRNA-mediated silencing in the germline (61), its preferential insertion at centromeres could provide an advantage for its continuous propagation since centromeres are typically not associated with heterochromatic marks (3, 27, 61, 62). Given the rapid evolution of centromere repeats and the lack of uniformity even within the five centromeres of *D. melanogaster*, targeting CENP-A chromatin preferentially represents an efficient way for *Jockey-3* to end up at centromeres. In turn, *Jockey-3* could benefit the host by promoting local transcription, which could facilitate chromatin remodeling during CENP-A deposition. Changes in expression for LINE1 modulate global chromatin accessibility during early mouse embryonic development, independently of both the LINE1 RNA or its protein products (63). Similarly, *Jockey-3* expression could promote local chromatin accessibility at centromeres. Future work will need to explore if the retention, the metaphase transcription of *Jockey-3*, or neither, are required for the integrity and maintenance of centromeric chromatin.

Global analyses of the chromatin-associated non-coding transcriptome in human embryonic stem cells showed that most RNA-DNA interactions are proximity based, with virtually none occurring in *trans*. Furthermore, TE-derived RNAs are frequently found associated with chromatin (64). Our results showing *cis* localization of *Jockey-3* are consistent with these findings. Even though we did not observe RNA-FISH signal in metaphase for the centromere-associated *Doc* retroelement, it is possible that additional centromere-derived RNAs contribute to the overall regulatory output of RNA-chromatin interactions at the centromere, similar to that proposed for genes (64).

Why *Jockey-3* RNAs are retained at centromeres remains unclear. RNA localization evidence does not differentiate between RNAs that are tethered to the centromere through the active transcriptional machinery from those complexed with centromeric proteins. These transcripts may simply be an incidental byproduct of the element’s transcription with no further regulatory role (45) or, like alpha-satellite RNAs, they could interact with centromeric proteins contributing to centromere integrity (9). Alternatively, transcript retention could serve as a mechanism for regulating *Jockey-3* transposition: it may function as an integral part of this retroelement’s mechanism of transposition or, conversely, as a defense strategy employed by genomes to prevent the transposon’s re-insertion in gene-encoding genomic regions.

*Jockey-3* transcripts form distinct, bright foci at metaphase centromeres, bearing similarity to RNA-rich nuclear condensates such as histone locus and Cajal bodies, or nucleoli (65). RNA has the ability to initiate condensate formation, supporting the nucleation of additional RNAs and proteins (66). In *S. pombe*, clustering of the centromeres by the Spindle Pole Body facilitates CENP-A assembly through this structure’s ability to attract high concentrations of CENP-A and its assembly factor (20). It is possible that high concentrations of *Jockey-3* transcripts produced in metaphase may aid in the maintenance of centromeres by attracting elevated levels of *Drosophila* CENP-A and its assembly factor CAL1 (48). This mechanism could depend more on the origin of the RNA (specifically, its derivation from centromeres) than its unique sequence.

## Supporting information

Table of Reagents

## Acknowledgements

We would like to thank Bo Reese from the UConn the Center for Genome Innovation in the Institute for Systems Genomics for assistance with sequencing and Vijender Singh (Computational Biology Core, Institute for Systems Genomics) and the UConn High Performance Computing for computational support. We are grateful to Jason Palladino and members of the Mellone lab for discussion and suggestions. This work was funded by NIH R35GM131868 to BGM. Additional support: NIH R01GM123312 to RJO, NSF MCB1844693 to AML, and NIH R35GM128857 to LC.

## Author contribution statement

Conceptualization: BM. Project administration: BM. Investigation: BS, RS, AA, OL, CC, LL. Formal analyses: BS, RS, AA, SJH, RD, BM. Visualization: BM, BS, RS, AA, SJH. Resources: BM, LC, AML. Methodology: All authors. Software: AA, SJH, RD. Validation: BM, BS, RS, AA, SJH. Supervision: BM, AML, LC, RJO. Writing-Original draft: BM, RS, SJH, BS, AA. Writing-Review and Editing: BM, RS, AA, SJH, BS, RJO. Funding Acquisition: BM, RJO, AML, LC.

## Methods

### *Drosophila* stocks and handling

Flies were reared on standard cornmeal, molasses, and yeast food (https://bdsc.indiana.edu) at 25°C, except for crosses for RNAi and sh-mediated knockdowns, which were carried out at 29°C. Experiments were performed in the following *D. melanogaster* stocks: laboratory stock iso-1 (Bloomington Drosophila Stock Center stock no. 2057: y1; Gr22b^iso-1^ Gr22d^iso-1^ cn^1^ CG33964^iso-1^ bw^1^ sp^1^; MstProx^iso-1^ GstD5^iso-1^ Rh6^1^); laboratory stock OreR (from A. Spradling lab); lacO (3^peri^, cytoband 80C4); UAS-CAL1-GFP-LacI and UAS-GFP-LacI maintained as heterozygous lines with the T(2;3)TSTL double balancer (43); sh-mCherry (Bloomington Drosophila Stock Center stock no. 35785) and sh-*Jockey-3*; gCID-EGFP-CENP-A/CID (P{gcid.EGFP.cid}III.2; (67). The GAL4 driver used was elav-GAL4 balanced with T(2;3)TSTL translocation balancer. The *D. simulans* stock used is w501 (gift of Andy Clark).

For all knockdowns, elav-GAL4 balanced with T(2;3)TSTL males were crossed with sh virgin females at 29°C. Non-tubby larvae, which carried both elav-GAL4 and the sh, were selected for dissections.

The sh-*Jockey-3* line was generated by PhiC31-mediated integration of pVALIUM20-sh-*Jockey3* at the attP2 landing site after injection by a commercial service (Best Gene). The *Jockey-3* hairpin was designed against the reverse-transcriptase region of *Jockey-3* using the DSIR website (http://biodev.extra.cea.fr/DSIR/DSIR.html), picking the one with the highest score. The sequences targeting *Jockey-3* were: 5’-ACGCTGGAACATCATGATCAA (Passenger strand) and 5’-TTGATCATGATGTTCCAGCGT (Guide strand). The oligos ordered included the passenger and guide strands flanked by standard flanking sequences. The resulting oligos were: 5’-ctagcagtACGCTGGAACATCATGATCAAtagttatattcaagcataTTGATCATGATGTTCCAGCGTgcg (Top strand) and 5’-aattcgcACGCTGGAACATCATGATCAAtatgcttgaatataactaACGCTGGAACATCATGATCAAactg (Bottom Strand). These top and bottom strands were annealed together creating overhangs and ligated into pVALIUM linearized with NheI and EcoRI.

### Cell culture

*Drosophila* Schneider (S2) cells were grown in Schneider’s media containing 10% FCS and anti-biotic/anti-mycotic mix at 25°C. Cells were passaged twice a week by diluting a cell resuspension to a million cells/ml.

### Stellaris probe design

Custom probes were designed using the Stellaris FISH probe designer. Probes were designed against the *Jockey-3* consensus sequence using ORF1 and ORF2 as targets. See Table of reagents for probes sequences.

### RNA extraction from brains and RT-qPCR

20-30 male larval brains were dissected in ice cold PBS DEPC and preserved in 150µl RNA later at -20°C. PBS DEPC was added to the brain suspension and spun to pellet the brains. The PBS/RNA later was removed and the brains were lysed in 300µl of TRIzol using a motorized pestle. RNA was extracted with Zymo Direct-zol RNA MiniPrep Kit (Cat#: 11-330) according to manufacturer’s instructions, except the in-column DNase I treatment was repeated twice. Samples were then treated with Turbo DNAse 2 to 3 times and then purified with the RNA Clean and Concentrator-5 Kit (Zymo Research Cat#: 11-325) according to the manufacturer’s instructions. cDNA was prepared with iScript Reverse Transcription Supermix following the manufacturer’s instructions. PCR was used to check cDNA quality and no DNA contamination in the no reverse transcriptase samples. qPCR was performed with iTaq Universal SYBR Green Supermix in 96 well plates, and ran on a BioRad qPCR thermocycler. Relative quantity was calculated with the Pfaffl method (68).

The PCR cycle was as follows: 95 °C 3 min for initial denaturation, then followed by 40 qPCR cycles. Each cycle has denaturation at 95 °C for 10s, annealing at 55°C for 20s and extension at 72°C for 20s.

### Primer design for targeting *Jockey-3*

We designed primers targeting the reverse-transcriptase domain within ORF2 from the *Jockey-3* consensus sequence using the Primer Design tool in Geneious Prime, avoiding the sequence targeted by the sh-*Jockey-3* itself. To determine which genomic copies are likely captured by these primers, we mapped the primers to the list *Jockey-3* insertions targeted by sh-*Jockey-3*, using the Map to Reference tool in Geneious Prime, allowing a maximum of 3 mismatches.

### Metaphase spread preparations from larval brains

All solutions were made up in DEPC milliQ water. Third instar larval brains were dissected (2-3 brains/slide) in PBS and all attached tissue and mouth parts were removed with forceps. Brains were immersed in 0.5% sodium citrate solution for 8 min in a spot well dish then moved to a 6µl drop of 45% acetic acid, 2% Formaldehyde on a siliconized (Rain X-treated) coverslip for 6 min. A poly-lysine coated glass slide was inverted and placed on the brains to make a sandwich. After flipping the slide and gently removing excess fixative between bibulous paper, the brains were squashed with the thumb by firmly pressing down. Slides were then immersed in liquid nitrogen and the coverslip was flipped off using a razor blade. Slides were then transferred to PBS for 5 min to rehydrate before proceeding with RNA-FISH/IF or IF/RNA-FISH. Monolayers brain preparation were performed using the same procedure except that acetic acid was omitted from the fixative.

### Mitotic spread preparations from S2 cells

3x10^5^ Schneider (S2) cells were collected in a tube for each slide and media was added to reach a volume of 475µl. The cells were treated for 1 hr with 0.5µg/ml colcemid (Sigma Aldrich) to induce mitotic arrest. Cells were then spun at 600g for 5 min in a centrifuge and resuspended in 250µl of 0.5% sodium citrate (DEPC treated) for 8 min. The cell suspension was loaded into a cytofunnel and spun for 5 min at 1200 rpm onto a poly-lysine coated slide using a cytocentrifuge (Shandon Cytospin 4, Thermo Fisher Scientific). The slides were immediately transferred to a coplin jar containing 100 ml of fixative (45% acetic acid and 2% formaldehyde in DEPC water) for 6 min. Slides were then washed 3 times with PBST (0.1% Triton) for 5 min while rocking at room temperature. Slides were stored in 70% ethanol at 4°C until IF/RNA-FISH.

### Mitotic spread preparations from ovaries

Ovary mitotic preparations were conducted as in (69). Mated adult females were anesthetized with CO2, then moved to a fresh 50 μL drop of PBS. Whole ovaries were dissected out and the carcass discarded. Using a needle, the tips of the ovaries were separated from later stages and immersed in 0.5% sodium citrate for 5 min, followed by fixation for 4 mins in 2 mL of fixative solution (45% acetic acid, 2.5% formaldehyde). Fixed tissues were moved to a 3 μL drop of 45% acetic acid on a siliconized coverslip (Rain X) and gently teased apart with a needle. A poly-L lysine coated glass slide was inverted onto the coverslip and pressed gently to spread the liquid to the edges of the coverslip. The slide and coverslip were squashed for 2 minutes using a hand clamp (Pony Jorgensen 32225), then immersed into liquid nitrogen for at least 5 minutes. Coverslips were immediately removed using a razor blade. The slide was then dehydrated by placing it in ice cold 70% ethanol for 2 hr at 4°C, and processed for RNA-FISH/IF.

### RNA-FISH/IF

Slides were immersed in PBST (0.1% Triton) and rocked for 10 min 3 times. Slides were transferred to 70% ethanol at 4°C overnight. Slides were rehydrated in PBST for 5 min and washed in wash buffer (2x SSC and 10% formamide) for 5 min while rocking. Without drying the brains, 50µl probe mix containing 45µl of Hybridization buffer (Stellaris), 5µl Formamide (10% formamide final) with 0.5µl of 12.5µM Stellaris smRNA FISH probes (0.125µM final concentration for Stellaris *Jockey-3* ORF1, ORF2, ORF2 antisense, *Doc*, *Rox1*). Brains were covered with a HybriSlip coverslip, sealed with rubber cement to prevent evaporation, and incubated at 37°C overnight in a humid chamber. Slides were then rinsed twice with wash buffer, washed twice in washing buffer for 30 min, and three times with 2X SSC for 10 min while gently shaking at RT. Slides were then post-fixed for 10 min in the dark in 100µl of 3.7% formaldehyde in PBS DEPC.

After 3 additional 5 min washes in PBST, the slides were then transferred to a coplin jar containing blocking buffer (1% BSA in PBST; PBS, 0.1% Triton-X) for 30 min while rocking. 50µl of primary antibodies (anti-CENP-C guinea pig polyclonal antibodies, 1:500) diluted in blocking buffer were applied to the slides, covered with parafilm and stored in a dark chamber at 4°C overnight. The following day, slides were washed 4 times with PBST for 5 min while rocking. Secondary antibodies (goat anti-guinea pig A488, 1:500) diluted in blocking buffer were applied to the brains, covered with a square of parafilm and incubated at room temperature for 1 hr. Slides were then washed 4 times in PBST for 5 min while rotating and again quickly in PBS for 3 min. Slides were mounted using SlowFade Gold containing 1µl/ml DAPI and a 22x22mm coverslip sealed with nail polish. The slides were stored in a dark environment to dry for 10 min before imaging.

### IF/RNA-FISH

Slides containing squashed larval brains were washed 3 times with PBST for 5 min on a rotator and transferred to 70% ethanol diluted at 4°C for 1 hr. Slides were then rehydrated for 5 min in PBST and processed for IF as described in the RNA-FISH/IF method above. After washing off the secondary antibodies, the slides were then processed for RNA-FISH without post-fixing, using Stellaris probes for *Jockey-3* and a lacO LNA probe Slides were mounted as described for RNA-FISH/IF.

### Sequential IF/RNA-FISH/DNA-FISH to detect lacO RNA at *de novo* centromeres

IF/RNA-FISH samples (anti-CENP-C guinea pig 1:500; lacO LNA, *Jockey-3* ORF2) were imaged and the list of points visited was saved. Coverslips were removed with a razor blade and the slides were washed in PBS for 10 min at room temperature while rocking. Slides were then washed three times with 4X SSC for 3 min, once with 2X SSCT for 5 min, and once with 50% formamide 2X SSC for 5 min at room temperature while rocking. 50 µl probe mix containing 13.5 µl 4X hybrid mix (8X SSC, 0.4% Tween20, 40% dextran sulfate, 34 µl formamide, 2µl RNase cocktail, 0.5 µl lacO LNA probe (100µM stock), 0.5 µl *dodeca* LNA probe (100µM stock) were added to the slide, covered with a hybrislip and sealed with rubber cement. Slides were incubated at 95°C for 5 min in a slide thermal cycler (Epperndorf) then transferred to a humid chamber and incubated at 37°C overnight in the dark. After incubation, the hybrislip and rubber cement were removed. Slides were then washed once at 37°C with 0.1X SSC for 10 min and twice at room temperature with 0.1X SSC for 10 min while rocking. Slowfade Gold containing DAPI was applied to the brains, covered with 22X40 mm or 22X22 mm coverslips, and sealed with nail polish. Imaging was performed by re-visiting the same point list.

### RNase treatments and quantification

For the RNase H treatments, male 3rd instar larval brain monolayers from a line expressing CENP-A/CID-EGFP under the control of the CENP-A/CID regulatory sequences (67) were processed for RNA-FISH using the *Jockey-3* ORF2 probe. Two slides were prepared. The following day, samples were imaged and point locations were recorded. Following imaging of these two pre-treatment slides, the coverslips were removed and the slides were briefly rinsed in PBS. RNase H treatment was performed with 10U of RNase H (cleaves the RNA when coupled with DNA; NEB) incubated for 2hr at 37°C in a dark humid chamber on one the slides, while the control slide was treated in the same way omitting the RNase H but including the buffer diluted in water. Slides were then washed once with PBS and mounted as described. The slides were then reimaged using the same settings as before, with the same points revisited. Quantification of the samples were done by counting the number foci of eCENP-A/CID-GFP and *Jockey-3* ORF1 probes within cells between the pretreatment and post-treatment. Values were plotted using Prism as a scatter plot. Statistical analysis was conducted using the t-test (unpaired).

For the RNase cocktail treatment, we generated male 3rd instar larval brain monolayers from eCID-GFP lines. Prior to RNA-FISH probe hybridization, 4U of RNase cocktail (RNase A and RNase T1, both targeting single-stranded RNA; Thermo Fisher) diluted in PBS was added to one slide (treated), while the other slide (untreated) only contained PBS. Samples were incubated at 37°C for 30 min. Samples were then washed for 5 min in PBS and hybridized with the *Jockey-3* ORF2 probe and Rox1 probes RNA-FISH. The following day the samples were imaged and point locations were recorded. Quantification of the samples was done by counting the number eCID-GFP and *Jockey-3* ORF2 foci within cells (N=100 cells) for both samples. Values were plotted as a scatter plot using Prism. Statistical analysis was conducted using the t-test (unpaired).

Our attempts to degrade the *Jockey-3* RNA-FISH signal from metaphase spreads with RNase H and RNase cocktail treatments were not successful, despite seeing *Rox1* signal become very weak or disappear. We hypothesize that the centromere/kinetochore protects *Jockey-3* RNA from degradation. We also performed these treatments after reversing the crosslinking at 80°C for 8 min as described in (21). However, heat treatment eliminated all *Jockey-3* RNA-FISH signal even in the absence of any RNase, precluding us from drawing any conclusions from these experiments.

### Imaging

All images were acquired at 25°C using an Inverted Deltavision ULTRA (Leica) equipped with a sCMOS pco.edge detector camera and with either a 100x/1.40 NA or 60x/1.42 NA oil objective using 0.2µm z-stacks. Mitotic spreads were imaged using the 100x objective. Tissue monolayers were imaged using either the 60x/1.42 NA or 100x/1.40 NA oil objectives. Image acquisition was performed using DeltaVision Ultra Image Acquisition software and image processing was performed using softWoRx software (Applied Precision). Images were deconvolved for 5 cycles using the conservative setting. All Stellaris probes for RNA-FISH were excited for 0.5s at 100% transmission for each z-slice image. Following deconvolution, images were quick-projected as maximum intensity projections using in-focus z-slices, a uniform scale was applied before saving images as Photoshop files. Images were minimally adjusted using Photoshop (Adobe) and assembled into figures in Illustrator (Adobe).

### Colocalization quantification for *Jockey-3* at centromeres

Metaphases were inspected in the CENP-C channel to identify centromeres and the presence of *Jockey-3* signal was determined by eye and recorded as colocalizing if present in at least one sister.

### Colocalization quantification for *Jockey-3* at *de novo* lacO centromeres

The presence of dicentrics causes chromosome breaks and rearrangements, making the identification of chromosomes difficult. Therefore, we selected metaphases with intact chromosome 3’s (identified with *dodeca* DNA-FISH) and with CENP-C signal at the 3peri location (identified with lacO DNA-FISH) for quantification. For the cis/trans *Jockey-3* ORF2 RNA quantification, the presence of *Jockey-3* RNA signal in the corresponding RNA-FISH images was determined by eye and recorded as present or absent. To determine if lacO transcripts were present, lacO RNA signal was determined by eye and recorded as present or absent. We selected metaphases with intact chromosome 3’s (identified with *dodeca* DNA-FISH) and with lacO at the 3peri location (identified with lacO DNA-FISH) for quantification.

### Fluorescence intensity quantifications

To measure *Jockey-3* signal at the centromeres of metaphase chromosomes, non-deconvolved in-focus z slices were quick-projected using the max intensity setting in SoftWorx. Polygons were drawn around the centromere of each chromosome using the edit polygons tool in the CENP-C channel then propagated to the *Jockey-3* channel to capture *Jockey-3* RNA max intensity fluorescence at the centromere. Similar polygons were used to capture background fluorescence for downstream calculations. Signal for sister centromeres were averaged and the average max intensity of the background fluorescence for that channel was subtracted. The measured max intensities for CENP-C and *Jockey-3* were plotted using Prism and compared.

For the quantification of metaphase spreads from sh-*Jockey-3* knockdowns, non-deconvolved 100x images were quick-projected in Softworks using the average intensity setting. Images were exported as TIFF and quantified with FIJI. In FIJI, a 400x400 pixel area including CENP-C, *Jockey-3* ORF1, and *Jockey-3* ORF2 foci on centromere Y was drawn to measure total intensities. Background intensities were set as lowest intensities in the square. Final fluorescence intensities in arbitrary units were calculated by subtracting background intensities from total intensities.

For the quantification of interphase spreads from sh-*Jockey-3* knockdowns, images were quick-projected in Softworks using the max intensity setting. Images were exported as TIFF and quantified with FIJI. In FIJI, entire nuclei were circled to measure raw max intensities of CENP-C, *Jockey-3* ORF1, and *Jockey-3* ORF2. Circles were then moved to the background area to measure background intensities. Final fluorescence intensities in arbitrary units were determined by subtracting background intensities from max intensities.

For the quantification of metaphase spreads from CAL1-GFP-LacI, *lacO* 3^peri^ and GFP-LacI, *lacO* 3^peri^, non-deconvolved 100x images were quick-projected in Softworks using the maximum intensity setting. Images were exported as TIFF and quantified with FIJI. In FIJI, a 400x400 pixel area including *lacO* foci on chromosome 3 was drawn to measure the total intensity. The background intensity was set as the average of 8 surrounding 400x400 pixel areas. The final fluorescence intensity in arbitrary units was calculated by subtracting the background intensity from the total intensity.

### Mapping *Jockey-3* RNA-FISH probes to centromeres

To determine how many probes are predicted to bind to each centromere, we mapped probes to the centromeric contigs extracted from the heterochromatin-enriched genome assembly from (24) using the map to reference tool in Geneious, using all default settings and allowing all best matches.

### Embryo collection, RNA extraction, and nuclei isolation for PRO-seq

Embryos were collected from 2-3 days old iso-1 flies at 25°C. Adult flies were kept in multiple cages on grape juice agar plates containing a small amount of fresh yeast paste. Collection plates from the first 1h were discarded and flies were allowed to lay embryos on grape juice agar plates for 12 hrs overnight. Embryos were rinsed thoroughly with water and egg wash (0.7% NaCl made in DEPC treated water plus 0.05% Triton-X 100) in a mesh basket. Embryos were then dechorionated with 50% bleach for 1 minute, rinsed thoroughly with tap water in a mesh basket, flash-frozen in liquid nitrogen, and stored at -80°C.

For RNA-seq, frozen embryos were resuspended in 300µl of TRI Reagent (Sigma Aldrich T9424) and homogenized using a motorized pestle. After centrifugation, RNA was extracted from the supernatant using the Zymo DirectZOL kit (Zymo Research) following the manufacturer’s instructions.

Embryo nuclei isolation was performed largely as described in (70). 50-100µl packed embryos were resuspended in 1mL cold buffer 1 (1M sucrose, 1M Tris pH 7.5, 1M MgCl2, 100% Triton X-100, 100mM EGTA, 1M DTT, 1x PTase inhibitor cocktail Roche, 20U/µl SUPERase In Ambion, 1M CaCl2), dounced in a 1m dounce homogenizer with a loose pestle 25 times, centrifuged at 900g for 2 min at 4°C to remove large debris, and dounced again with a tight pestle 15 times on ice. Nuclei were pelleted at 800g for 10 min at 4°C and washed twice in buffer 1 and once in freezing buffer (1M Tris pH 8, 100% glycerol, 100mM MgAc2, 0.5M EDTA, 1M DTT, 1x PTase inhibitor cocktail Roche, 20U/µl SUPERase In Ambion). Nuclei were resuspended in freezing buffer, flash-frozen, and stored at -80°C until use.

### Nuclei and RNA isolation from larval brains for PRO-seq and RNA-seq

Wandering larvae (3rd instar; OreR stock for PRO-seq and iso-1 for RNA-seq) were washed and dissected in PBS. Approximately 125 brains were dissected, flash frozen in liquid nitrogen, and stored at -80°C. Nuclei isolation was performed as described for the embryos but using a 0.5ml dounce homogenizer. Total RNA extraction was performed as described for embryos.

### PRO-seq library generation, pre-processing and alignment

PRO-seq libraries were prepared as previously described (40). 0.9-4.5 x 10^6^ nuclei were mixed with permeabilized 1 x 10^6^ Hela nuclei (as spike-in) in 4-biotin-NTP run-on reactions. Run-on RNA was then base-hydrolyzed for 20 min on ice and enriched using M280 streptavidin beads and TRIzol extraction. After amplification, libraries were purified by polyacrylamide gel electrophoresis (PAGE) to remove adapter-dimers and to select molecules below 650 bp in size. Libraries were then sequenced on an Illumina NextSeq 500/550, producing paired-end 100bp reads. We obtained approximately 71 million reads (0-12h embryos) and 55 million reads (L3 brains).

Raw fastq files were first trimmed for quality (q 20), length (20 bp), and adapter sequences removed using cutadapt (71). For use with Bowtie 2 (72), paired-end reads were aligned to a combined Human (GRCh38) - *Drosophila* heterochromatin-enriched assembly (24) using default “best match” parameters. A position sorted bam file containing reads mapping to *Drosophila* was de-duplicated (removal of duplicate reads) using Picard’s MarkDuplicates (http://broadinstitute.github.io/picard/). It should be noted that read duplicates can emerge during library preparation via PCR, but in the case of PRO-seq they can also be the result of RNA polymerase pausing; since we cannot be sure which is the case with this method, we opted to remove duplicate reads to be conservative. This de-duplicated bam was then processed into a bed file using BEDtools (Quinlan and Hall, 2010), which was used for generation of a 3 ’end only (RNA polymerase occupancy position) bed file. This 3 ’end only bed file was then used for either: 1) counting read abundance and coverage with BEDtools, or 2) BigWig file generation for visualization in the Integrated Genome Viewer (IGV) (Robinson et al., 2011).

For use with Bowtie, read 1 was reverse-complemented using the fastx-toolkit (http://hannonlab.cshl.edu/fastx_toolkit) and then aligned to a combined Human (GRCh38) - *Drosophila* heterochromatin-enriched assembly using k-100 parameters (reporting up to 100 mapped loci for each read). Since the purpose of this mapping method was to include multi-mappers as a representation of the “upper bounds” of transcription, de-duplication was not performed on the k-100 read set. Sorted bam files containing reads mapping to *Drosophila* were processed into bed files using BEDtools (73), which were used for either: 1) unique 21-mer filtering (described below in “Meryl unique k-mer filtering”), or 2) generation of 3 ’end only (RNA polymerase occupancy position) bed files. In the case of option 2) these 3 ’end only bed files were then use for either: 1) counting read abundance and coverage with BEDtools, or 2) BigWig file generation for visualization in the Integrated Genome Viewer (IGV) (74).

### RNA-seq library generation, pre-processing, and alignment

RNA-seq libraries were generated using 200ng of RNA from 0-12h embryos or 3rd instar larval brains using Illumina stranded total RNA prep, with the ligation performed with Ribo-Zero Plus and sequenced on Illumina TruSeq Stranded total RNA library prep kit, producing 150bp paired-end reads. We obtained approximately 46 million reads (0-12h embryos) and 33 million reads (L3 brains).

Raw fastq files were first trimmed for quality (q 20) and length (100 bp), and then adapter sequences removed using cutadapt (71) before being aligned to a *Drosophila* heterochromatin-enriched assembly (24) as paired-end reads using either Bowtie 2 (72) default “best match” parameters or Bowtie k-100 (75). HeLa spike-ins were not included in RNA-seq data and therefore, did not need to be removed. In each case, sorted bam files were processed into bed files using BEDtools (73), which were used for one of the following: 1) unique 51-mer filtering, 2) counting read abundance and coverage with BEDtools, or 3) BigWig file generation (BEDtools, GenomeBrowser/20180626) for visualization in the Integrated Genome Viewer (IGV) (74).

### Meryl unique k-mer filtering

Single copy k-mers were generated from *Drosophila* heterochromatin-enriched assembly using Meryl (76). We chose the length of single-copy k-mers (21 versus 51-mers) to use for filtering based on the length of the library insert, which is smaller for PRO-seq than for RNA-seq. Bed files of the mapped reads were used to filter through Meryl single copy k-mers using overlapSelect with the option ‘-overlapBases=XXbp ’(XX represents the length of the single copy k-mers (21-mer or 51-mer); GenomeBrowser/20180626). This locus-level filtering requires a minimum of the entire length of k-mer should overlap with a given read in order to be retained. The bed files from all RNA-seq mapping methods (default, k-100, and k-100 51-mer filtered) were used for read counts for repeats and BigWig file generation of IGV visualization (74). The bed files from all PRO-seq mapping methods (default, k-100, and k-100 21-mer filtered) were first processed into 3 ’end only (RNA polymerase occupancy position) bed files before being used for read counts across repeats and BigWig file generation for IGV visualization.

### Centromere heat maps for PRO-seq and RNA-seq data

The density of all centromeric repeats was obtained by counting the number of reads mapping to each repeat and dividing it by the number of total reads mapping to that centromeric contig. Read counts of all repeats were obtained with bedtools coverage -counts option. All heatmaps were generated with the ggplot2 R package.

### CUT&Tag from embryos

2-12h old *Drosophila* iso-1 embryos were collected from cages containing grape-juice agar plates with yeast paste incubated overnight at 25°C. Embryos were washed in embryo wash buffer (0.7% NaCl, 0.04% Triton-X100) and then were dechorionated with 50% bleach for 30s. Embryos were lysed in 1ml buffer B (pH7.5, 15mM Tris-HCl, 15mM NaCl, 60mM KCl, 0.34M Sucrose, 0.5mM Spermidine, 0.1% β-mercaptoethanol, 0.25mM PMSF, 2mM EDTA, 0.5mM EGTA) using a homogenizer and filtered through a mesh to remove large debris. Nuclei were spun at 5000g for 5 min and resuspended in 500µl of buffer A (pH7.5, 15mM Tris-HCl, 15mM NaCl, 60mM KCl, 0.34M Sucrose, 0.5mM Spermidine, 0.1% β-mercaptoethanol, 0.25mM PMSF) twice. The final pellet was resuspended in CUT&Tag wash buffer (20mM HEPES pH 7.5, 150mM NaCl, 0.5 mM Spermidine) to a final concentration of 1 million nuclei/ml.

CUT&Tag was performed on approximately 50,000 nuclei per sample using the pA-Tn5 enzyme from Epycpher, following the manufacturer’s instructions (CUT&Tag Protocol v1.5; (46). We used a rabbit anti-Cid/CENP-A antibody (Active Motif cat. 39713, 1:50) and rabbit anti-IgG as negative control (1:100). For the library preparation, we used the primers from (77). Before final sequencing, we pooled 2µl of each library and performed a MiSeq run. We used the number of resulting reads from each library to estimate the relative concentration of each library and ensure an equal representation of each library in the final pool for sequencing. We sequenced the libraries in 150-bp paired-end mode on HiSeq Illumina. We obtained around 6-9 million reads per library, except for the IgG negative control which typically yields much lower reads.

### CUT&Tag mapping

Raw fastq files of CUT&Tag data were trimmed using trimgalore with these options --paired --nextera -- length 35 --phred33 and read quality was assessed with FASTQC. Reads were mapped to Drosophila heterochromatin-enriched assembly with Bowtie2. And MACS2 callpeak was used to call peaks using the IgG as our input control (options -c IgG.bam -f BAMPE -g dm -q 0.01 -B --callsummits). The CENP-A domains were defined based on MACS2 peaks and deepTools bamCompare (78) read coverage. The CENP-A domain for each centromere was determined from the first to the last MACS2 peak. Non-centromeric CENP-A domains were defined based on MACS2 peaks alone without having a single domain for each contig as compared to centromeres. As per Fig. 5B, MACS2 signal intensity values were averaged (BEDtools map -o mean; (73)) from the narrowPeak file across each CENP-A domain.

### Statistical tests

All *Jockey-3* sequences were extracted from *Drosophila* heterochromatin-enriched assembly annotations using BEDtools (73) and labeled as CENP-A-CEN, CENP-A-nonCEN, or nonCENP-A (requiring at least 1bp overlap with MACS2 CENP-A domains) using BEDtools map -o collapse. *Jockey-3* copies were also labeled as either full-length (FL; if containing a full ORF2) or truncated. Lastly, *Jockey-3* copies were categorized by age based on their divergence from the *Jockey-3* consensus sequence from (61), wherein less than 1% divergence was categorized as ‘young’ and greater than or equal to 1% was categorized as ‘old’ (61). It should be noted that the age categorization from Hemmer et al. (61)was available for 326 out of the 329 copies included in all our other analyses. PRO-seq read counts were obtained with BEDtools coverage -counts (requiring at least 1bp overlap) for all *Jockey-3* copies in the genome, as well as for each CENP-A domain and CENP-A-nonCEN-sized random interval. Unique 21-mer coverage per *Jockey-3*, as well as *Jockey-3* coverage per CENP-A domain was assessed using BEDtools coverage. Unpaired *t* tests were performed to quantify differences and determine significance. Scatter box plots and bar graphs were generated via GraphPad Prism (v10.1.1). Heatmaps representing PRO-seq transcriptional profiles were generated with deepTools computeMatrix and plotHeatmap (Ramirez et al., 2016). Specific plotting parameters include: --averageTypeBins max, --averageTypeSummaryPlot mean, and --zMax 9.

### Code and data access

All code for analyses and figures are available on Github https://github.com/bmellone/Dmel-Centromere-Transcription. All sequencing data is available on NCBI under Bioproject PRJNA1082342.

## Supplemental Figure Legends

**Figure S1: PRO-seq reads aligned to genes show expected enrichment of RNA polymerase occupancy at gene promoters.** Heatmaps of RNA polymerase occupancy mapped using Bowtie 2 default “best match” for antisense (blue) and sense (red) strands per gene. Composite profiles (line graphs) across all genes are shown along the top.

**A** All genes are anchored to the 5 ’end (transcription start site (TSS)) with a specified distance into the gene body denoted in the bottom right (2.5kb), and a specified distance away from the gene body denoted in the bottom left (0.5kb). The dotted line per heatmap denotes the static end of each gene as they are sorted longest to shortest from top to bottom. This highlights the anticipated enrichment of RNA polymerase at the promoter.

**B** All genes are scaled to the same size with a specified distance on either side of the gene body denoted in the bottom corners (0.1kb). The genes are included in the heatmap based on transcriptional signal intensity from top to bottom. This highlights RNA polymerase activity across the entire gene, with an enrichment at the promoter, reduction over the gene body, and a slight enrichment at the 3 ’end indicative of polymerase slow-down as termination occurs.

**C** Same as shown in **A**, except using PRO-seq reads that have been deduplicated. This highlights the overall preservation of the transcriptional pattern following deduplication with an expected loss of reads, predominantly at the promoter since this is where polymerase density naturally highest, meaning duplicate reads are more likely at this position.

**D** Same as shown in **B**, except using PRO-seq reads that have been deduplicated. This highlights the overall preservation of the transcriptional pattern following deduplication with an expected loss of reads at the promoter as well as at the 3 ’end where termination is occurring.

**Fig. S2:FL vs truncated k-100 and k-100 filtered PRO-seq**

**A** PRO-seq read density scatter boxplot comparisons between full-length (FL) and truncated *Jockey-3* copies, regardless of genome location. Mapping was done with Bowtie k-100 and k-100 21-mer filtered using single-end reads. Unpaired t-tests (Student’s t-test) were performed indicating a significant difference (****, p < 0.0001) between each group illustrating a consistent trend seen across all three mapping methods (Fig. 1B). Standard deviation error bars are shown.

**B** Meryl unique 21-mer coverage for FL and truncated *Jockey-3* copies. An unpaired t test (Student’s t-test) was performed indicating a significant difference (****, p < 0.0001), wherein truncated copies have more unique 21-mers as a result of having accumulated more mutations over time making them less similar to each other.

**Fig. S3:PRO-seq and RNA-seq from larval brains**. PRO-seq, RNA-seq signals for 3rd instar larval brains across all *D. melanogaster* centromeres. Top track shows sense, bottom, antisense. Tracks show read coverage with three mapping methods: Bowtie 2 default best match (“lower bounds”; yellow), over-fit (“upper bounds”; gray) and a filtered over-fit (“medium bounds”; blue). For PRO-seq we Bowtie1 k-100 for over-fit, and Bowtie1 k-100 21-mer filtered for medium bounds. For RNA-seq we used Bowtie2 k-100 for over-fit and Bowtie2 k-100 51-mer filtered for medium bounds. Repeat annotation is shown on top (see legend for details), with unique 21 and 51-mers (black) used for the filtering shown below. The k-mer tracks illustrate the regions that lack sequence specificity and are therefore most prone to read loss through k-mer filtering. Coordinates shown are kilobases. Dotted red line indicates the boundaries of the islands.

**Figure S4:RNA-FISH detects the chromosome-associated non-coding RNA *Rox1***. RNA-FISH/IF on *D. melanogaster* (iso-1) mitotic chromosomes from male larval brains with the ORF2 of *Jockey-3* probe (yellow), a *Rox1* probe (cyan), and with CENP-C antibodies (green). DNA is stained with DAPI (magenta). Bars 1µm. Arrow points to *Rox1* (yellow) localization on the arms of the X chromosome.

**Figure S5:*Jockey-3* RNA localization on centromeres of individual chromosomes**. RNA-FISH/IF on *D. melanogaster* (iso-1) mitotic chromosomes from male larval brains. Individual chromosomes showing centromeric signal for each *Jockey-3* probe set (ORF2, ORF2 anti, and ORF1) (yellow) on chromosomes (X, Y, 2, 3, and 4), CENP-C showing the centromere (green), and DAPI (magenta). Insets show CENP-C and *Jockey-3* signals. Bars 1µm

**Figure S6 *Jockey-3* RNA localizes to mitotic centromeres in other tissues and in *D. simulans***

**A** RNA-FISH/IF on mitotic chromosomes from *D. melanogaster* (iso-1) adult ovaries. IF for CENP-C (green) and RNA-FISH for *Jockey-3* ORF2 (yellow). DNA is stained with DAPI (magenta).

**B** RNA-FISH/IF on mitotic spreads from S2 cells. IF for CENP-C (green), and RNA-FISH for *Jockey-3* ORF2 (yellow) and SatIII (found on X and 3rd chromosomes; cyan). DNA is stained with DAPI (magenta).

**C** RNA-FISH/IF on *D. simulans* (laboratory stock w501) mitotic chromosomes from male larval brains. IF with CENP-C (green) and RNA-FISH for *Jockey-3* ORF2 and *Rox1* (stains the X, control; cyan). DNA is stained with DAPI (magenta). Bar 1µm

**Figure S7:RNA foci detected with *Jockey-3* ORF2 correspond to RNA, not DNA**

**A** RNA-FISH on *D. melanogaster* (iso-1) mitotic chromosomes from male larval brains for the 3 ’sense of *Jockey-3* (yellow) and an OligoPaint for 61C7 (green). Green arrow indicates presence of signal, white arrow indicates lack of signal.

**B** DNA-FISH on *D. melanogaster* (iso-1) mitotic chromosomes from male larval brains with stellaris FISH probes for ORF2 of *Jockey-3* (yellow) and an OligoPaint for 61C7 (green). Green arrow indicates presence of signal, white arrow indicates lack of signal. Bars 5µm.

**Figure S8:RNase treatments result in a decrease in *Jockey-3* RNA-FISH signal**

**A** RNA-FISH with *Jockey-3* ORF2 on *D. melanogaster* male larval brain monolayers expressing eCENP-A-EGFP (green). DNA is stained with DAPI (magenta). Shown are the before and after treatments with and without RNase H.

**B** Quantification of the number of ORF2 *Jockey-3* foci before and after RNase H treatment (not significant for before and p<0.0001 for after treatment). Quantification of the number of eCENP-A-EGFP foci before and after RNase H treatment (p=0.367 for before and not significant for after treatment). N=1 brain, n=107 cells quantified for before treatment and n=86 cells for after treatment).

**C** RNA-FISH with *Jockey-3* ORF2 and *Rox1* (cyan; control) on *D. melanogaster* male larval brain monolayers expressing eCENP-A-EGFP (under the endogenous CENP-A promoter; green).

**D** Quantification of eCENP-A-GFP (N=1 brain, n=100 cells; *p = 0.0292) and ORF2 *Jockey-3* foci (N=1 brain, n=100 cells; ****p = <0.0001).

**Figure S9:Quantification of non-centromeric *Jockey-3* foci in mitotic cells.** Graph showing the non-centromeric localization of *Jockey-3* (ORF2, ORF2 anti, and ORF1) on mitotic chromosomes from larval brain squashes. XR, 4L, and 4R were not quantified since these arms are cytologically too small and too close to the centromeres to be distinguished. ORF2 (N=3 brains, n=83 spreads), ORF2 anti (N=3 brains, n=28 spreads), and ORF1 (N=4 brains, n=69 spreads).

**Figure S10:RNA-FISH *Jockey-3* foci are present during interphase**

**A** RNA-FISH/IF with ORF2 (yellow) and ORF1 (magenta) *Jockey-3* probes and CENP-C (green) on interphase cells from male larval brain squashes.

**B** IF/RNA-FISH as in A on S2 cells (30% of cells have at least 1 co-localizing CENP-C/*Jockey-3* spot, n=54; note that S2 cells do not have a Y chromosome).

**C** RNA-FISH/IF as in A on ovary squashes. Insets show magnification of centromeres in the box.

**D** Graph showing the average number of *Jockey-3* ORF2 foci that co-localize with CENP-C (cen) versus not (non-cen).|

**E** Graph of the average number of centromeric *Jockey-3* foci in interphase versus metaphase cells. N=3 brains, n=30-105 cells per brain.

**F** Graph showing the % of cells showing 2 or more *Jockey-3* foci co-localizing with CENP-C in interphase versus mitosis. Data in **D-F** is all from the same 3 male larval brains as in **A**.

**Figure S11:RNA-FISH for centromeric retroelement *Doc***. RNA-FISH/IF on *D. melanogaster* (iso-1) male larval brain squashes. Immunofluorescence for CENP-C (green), and RNA-FISH for *Jockey-3* ORF2 (yellow) and *DOC* sense (cyan). *Doc* is present in the islands of centromere X and 4. Dashed box shows the X centromere lacking *Doc* signal. The solid line box shows a centromere with *Doc* signal in interphase. DNA is stained with DAPI (magenta).

**Figure S12:IGV tracks for CENP-A CUT&Tag**. IGV tracks showing CUT&Tag signals for 0-12h embryos across all *D. melanogaster* centromeres. Top track shows color-coded repeat annotation (details in legend). CUT&Tag track shows CENP-A enrichment in gray. Red dotted line shows the span of the CENP-A domain we defined for each centromere. Predicted MACS2 peaks for CUT&Tag data are shown in bottom track (black).

**Figure S13:CENP-A associated k-100 and k-100 filtered PRO-seq**.

**A** Bar graph illustrating the proportion of CENP-A associated *Jockey-3* copies expressed within the centromere and outside the centromere. PRO-seq mapping was done with Bowtie k-100 and k-100 21-mer filtered using single-end reads. Expression is defined as having at least two PRO-seq read overlaps. The trend difference seen between Bowtie 2 default and Bowtie k-100 methods can be attributed to the lower unique 21-mer coverage of CENP-A copies allowing more reads to map to these copies.

**B** Meryl unique 21-mer coverage for CENP-A associated *Jockey-3* copies based on centromeric loci designation. Unpaired t tests (Student’s t-test) were performed indicating a significant difference (****, p < 0.0001; ***, p < 0.001; ns (non-significant), p > 0.05) between each group. Standard deviation error bars are shown.

## Supplemental Table Legends

**Table S1:PRO-seq read and unique 21-mer coverage across all *Jockey-3* loci**. Table showing all 329 *Jockey-3* copies per CENP-A and centromeric association further distinguished by age based on divergence from consensus (<1%). PRO-seq read coverage for all three mapping methods are included: Bowtie 2 default “best match” using paired-end reads (post-deduplication), and Bowtie k-100 and Bowtie k-100 21-mer filtered, both using single-end reads. Coverage of Meryl unique 21-mers per copy is also shown. Data included was used for **Figs. 1B-C**, **Figs. 5F-G, Figs. S2 and S13, and Fig. 6**. Note: This table includes three truncated, old nonCENP-A copies indicated by an asterisk (*) in columns E & F, which are included in all analyses except those represented in **Fig. 6**.

**Table S2:Read counts for heatmaps**. Table showing the PRO-seq read count for each centromeric repeat within all centromere contigs. This data was used to generate the heatmaps shown in **Fig. 1**.

**Table S3:*Jockey-3* RNA-FISH probe sequences mapped across the genome.** The table shows the chromosome, contig, and coordinates of every *Jockey-3* copy in the genome. The first tab shows just the full-length copies, the second shows all the centromeric and the last all non-centromeric insertions. Indicated are the type of chromatin they are found in (if known; designated as in (24)), approximate cytological location and number of probes predicted to bind. This information was used for the graph in **Fig. 2F**.

**Table S4:CENP-A domain loci, both centromeric and non-centromeric.** Table showing all five centromeric and 333 non-centromeric CENP-A domains as defined by MACS2 peak calls from CUT&Tag data. Size (basepairs), average MACS2 peak signal intensity, and PRO-seq read overlap is shown per CENP-A domain. PRO-seq mapping was done with Bowtie 2 default “best match” using paired-end reads, post-deduplication. Data included was used for **Figs. 5A-B**.

**Table S5:Proportion of *Jockey-3* copies expressed based on PRO-seq read overlap.**

**A** Table showing the number of *Jockey-3* copies (FL and truncated) expressed per CENP-A and centromeric association. Expression is defined as having at least two PRO-seq read overlaps. All three mapping methods are included: Bowtie 2 default “best match” using paired-end reads (post-deduplication), and Bowtie k-100 and Bowtie k-100 21-mer filtered, both using single-end reads. Data included (representing 329 copies) was used for **Figs. 5C-D and Fig. S13**.

**B** Same as shown in **A**, except further distinguished by age based on divergence from consensus (<1%) and only representing the 326/329 copies with age distinctions (young vs. old). Data included was used for **Fig. 6**.

**Table S6:Summary of CENP-A-associated truncated and full-length (FL) *Jockey-3* insertions.** Table showing the distribution of all 329 *Jockey-3* copies associated with CENP-A and/or centromeres across the genome. A column for other repeats, excluding *Jockey-3*, is shown to emphasize the enrichment of *Jockey-3* associated with CENP-A. Note: this list does include 3 truncated, old nonCENP-A copies indicated by an asterisk (*), which are include in all analyses except those represented in Figure 6. Data included was used for **Fig. 5E**.

## Notes

### Competing Interest Statement

The authors have declared no competing interest.

### Summary of Updates

New data added (Fig. 6). Text and title have also been updated.

## Bibliography

1. Palmer DK, O’Day K, Wener MH, Andrews BS, Margolis RL. A 17-kD centromere protein (CENP-A) copurifies with nucleosome core particles and with histones. J Cell Biol. 1987;104(4):805–15.

2. Heterochromatic deposition of centromeric histone H3-like proteins, (2000).

3. McKinley KL, Cheeseman IM. The molecular basis for centromere identity and function. Nat Rev Mol Cell Biol. 2016;17(1):16–29.

4. Marshall OJ, Chueh AC, Wong LH, Choo KH. Neocentromeres: new insights into centromere structure, disease development, and karyotype evolution. Am J Hum Genet. 2008;82(2):261–82.

5. Ventura M, Antonacci F, Cardone MF, Stanyon R, D’Addabbo P, Cellamare A, et al. Evolutionary formation of new centromeres in macaque. Science. 2007;316(5822):243–6.

6. Ohkuni K, Kitagawa K. Endogenous transcription at the centromere facilitates centromere activity in budding yeast. Curr Biol. 2011;21(20):1695–703.

7. Hedouin S, Logsdon GA, Underwood JG, Biggins S. A transcriptional roadblock protects yeast centromeres. Nucleic Acids Res. 2022;50(14):7801–15.

8. Quenet D, Dalal Y. A long non-coding RNA is required for targeting centromeric protein A to the human centromere. eLife. 2014;3:e03254.

9. McNulty SM, Sullivan LL, Sullivan BA. Human Centromeres Produce Chromosome-Specific and Array-Specific Alpha Satellite Transcripts that Are Complexed with CENP-A and CENP-C. Dev Cell. 2017;42(3):226–40 e6.

10. Bury L, Moodie B, Ly J, McKay LS, Miga KH, Cheeseman IM. Alpha-satellite RNA transcripts are repressed by centromere-nucleolus associations. eLife. 2020;9.

11. Hoyt SJ, Storer JM, Hartley GA, Grady PGS, Gershman A, de Lima LG, et al. From telomere to telomere: The transcriptional and epigenetic state of human repeat elements. Science. 2022;376(6588):eabk3112.

12. Blower MD. Centromeric Transcription Regulates Aurora-B Localization and Activation. Cell Rep. 2016;15(8):1624–33.

13. Grenfell AW, Heald R, Strzelecka M. Mitotic noncoding RNA processing promotes kinetochore and spindle assembly in Xenopus. J Cell Biol. 2016;214(2):133–41.

14. Topp CN, Zhong CX, Dawe RK. Centromere-encoded RNAs are integral components of the maize kinetochore. Proc Natl Acad Sci U S A. 2004;101(45):15986–91.

15. Carone DM, Zhang C, Hall LE, Obergfell C, Carone BR, O’Neill MJ, et al. Hypermorphic expression of centromeric retroelement-encoded small RNAs impairs CENP-A loading. Chromosome Res. 2013;21(1):49–62.

16. Chen CC, Bowers S, Lipinszki Z, Palladino J, Trusiak S, Bettini E, et al. Establishment of Centromeric Chromatin by the CENP-A Assembly Factor CAL1 Requires FACT-Mediated Transcription. Dev Cell. 2015;34(1):73–84.

17. Chueh AC, Northrop EL, Brettingham-Moore KH, Choo KH, Wong LH. LINE retrotransposon RNA is an essential structural and functional epigenetic component of a core neocentromeric chromatin. PLoS Genet. 2009;5(1):e1000354.

18. Murillo-Pineda M, Valente LP, Dumont M, Mata JF, Fachinetti D, Jansen LET. Induction of spontaneous human neocentromere formation and long-term maturation. J Cell Biol. 2021;220(3).

19. Naughton C, Huidobro C, Catacchio CR, Buckle A, Grimes GR, Nozawa RS, et al. Human centromere repositioning activates transcription and opens chromatin fibre structure. Nat Commun. 2022;13(1):5609.

20. Wu W, McHugh T, Kelly DA, Pidoux AL, Allshire RC. Establishment of centromere identity is dependent on nuclear spatial organization. Curr Biol. 2022;32(14):3121–36 e6.

21. Bobkov GOM, Gilbert N, Heun P. Centromere transcription allows CENP-A to transit from chromatin association to stable incorporation. J Cell Biol. 2018.

22. Cardinale S, Bergmann JH, Kelly D, Nakano M, Valdivia MM, Kimura H, et al. Hierarchical inactivation of a synthetic human kinetochore by a chromatin modifier. Mol Biol Cell. 2009;20(19):4194–204.

23. Rosic S, Kohler F, Erhardt S. Repetitive centromeric satellite RNA is essential for kinetochore formation and cell division. J Cell Biol. 2014;207(3):335–49.

24. Chang CH, Chavan A, Palladino J, Wei X, Martins NMC, Santinello B, et al. Islands of retroelements are major components of Drosophila centromeres. PLoS Biol. 2019;17(5):e3000241.

25. Garrigan D, Kingan SB, Geneva AJ, Andolfatto P, Clark AG, Thornton KR, et al. Genome sequencing reveals complex speciation in the Drosophila simulans clade. Genome Res. 2012;22(8):1499–511.

26. Jagannathan M, Warsinger-Pepe N, Watase GJ, Yamashita YM. Comparative Analysis of Satellite DNA in the Drosophila melanogaster Species Complex. G3 (Bethesda). 2017;7(2):693–704.

27. Courret C, Hemmer L, Wei X, Patel PD, Santinello B, Geng X, et al. Rapid turnover of centromeric DNA reveals signatures of genetic conflict in Drosophila. BioRXiv. 2023.

28. Presting GG. Centromeric retrotransposons and centromere function. Curr Opin Genet Dev. 2018;49:79–84.

29. Choi ES, Stralfors A, Castillo AG, Durand-Dubief M, Ekwall K, Allshire RC. Identification of noncoding transcripts from within CENP-A chromatin at fission yeast centromeres. J Biol Chem. 2011;286(26):23600–7.

30. Chan FL, Marshall OJ, Saffery R, Kim BW, Earle E, Choo KH, et al. Active transcription and essential role of RNA polymerase II at the centromere during mitosis. Proc Natl Acad Sci U S A. 2012;109(6):1979–84.

31. Chan FL, Wong LH. Transcription in the maintenance of centromere chromatin identity. Nucleic Acids Res. 2012;40(22):11178–88.

32. Choi ES, Stralfors A, Catania S, Castillo AG, Svensson JP, Pidoux AL, et al. Factors that promote H3 chromatin integrity during transcription prevent promiscuous deposition of CENP-A(Cnp1) in fission yeast. PLoS Genet. 2012;8(9):e1002985.

33. Mellone BG, Fachinetti D. Diverse mechanisms of centromere specification. Curr Biol. 2021;31(22):R1491–R504.

34. Klein SJ, O’Neill RJ. Transposable elements: genome innovation, chromosome diversity, and centromere conflict. Chromosome Res. 2018;26(1-2):5–23.

35. Rosic S, Erhardt S. No longer a nuisance: long non-coding RNAs join CENP-A in epigenetic centromere regulation. Cell Mol Life Sci. 2016;73(7):1387–98.

36. Grenfell AW, Strzelecka M, Heald R. Transcription brings the complex(ity) to the centromere. Cell Cycle. 2017;16(3):235–6.

37. Perea-Resa C, Blower MD. Centromere Biology: Transcription Goes on Stage. Mol Cell Biol. 2018;38(18).

38. Mills WK, Lee YCG, Kochendoerfer AM, Dunleavy EM, Karpen GH. RNA from a simple-tandem repeat is required for sperm maturation and male fertility in Drosophila melanogaster. eLife. 2019;8.

39. Wei X, Eickbush DG, Speece I, Larracuente AM. Heterochromatin-dependent transcription of satellite DNAs in the Drosophila melanogaster female germline. eLife. 2021;10.

40. Mahat DB, Kwak H, Booth GT, Jonkers IH, Danko CG, Patel RK, et al. Base-pair-resolution genome-wide mapping of active RNA polymerases using precision nuclear run-on (PRO-seq). Nat Protoc. 2016;11(8):1455–76.

41. Hemmer NS, Geng X, Courret C, Navarro-Domínguez B, Speece I, Wei X, Altidor E, Chaffer J, Sproul J, Larracuente A. Centromere-associated retroelement evolution in Drosophila melanogaster reveals an underlying conflict. BioRXiv. 2023.

42. Meller VH, Wu KH, Roman G, Kuroda MI, Davis RL. roX1 RNA paints the X chromosome of male Drosophila and is regulated by the dosage compensation system. Cell. 1997;88(4):445–57.

43. Palladino J, Chavan A, Sposato A, Mason TD, Mellone BG. Targeted De Novo Centromere Formation in Drosophila Reveals Plasticity and Maintenance Potential of CENP-A Chromatin. Dev Cell. 2020;52(3):379–94 e7.

44. McNulty SM, Sullivan BA. Centromere Silencing Mechanisms. Prog Mol Subcell Biol. 2017;56:233–55.

45. Bassett AR, Akhtar A, Barlow DP, Bird AP, Brockdorff N, Duboule D, et al. Considerations when investigating lncRNA function in vivo. Elife. 2014;3:e03058.

46. Kaya-Okur HS, Wu SJ, Codomo CA, Pledger ES, Bryson TD, Henikoff JG, et al. CUT&Tag for efficient epigenomic profiling of small samples and single cells. Nat Commun. 2019;10(1):1930.

47. Nechemia-Arbely Y, Miga KH, Shoshani O, Aslanian A, McMahon MA, Lee AY, et al. DNA replication acts as an error correction mechanism to maintain centromere identity by restricting CENP-A to centromeres. Nat Cell Biol. 2019;21(6):743–54.

48. Chen CC, Dechassa ML, Bettini E, Ledoux MB, Belisario C, Heun P, et al. CAL1 is the Drosophila CENP-A assembly factor. J Cell Biol. 2014;204(3):313–29.

49. Murillo-Pineda M, Jansen LET. Genetics, epigenetics and back again: Lessons learned from neocentromeres. Exp Cell Res. 2020;389(2):111909.

50. Propagation of centromeric chromatin requires exit from mitosis, (2007).

51. Incorporation of Drosophila CID/CENP-A and CENP-C into centromeres during early embryonic anaphase, (2007).

52. Mellone BG, Grive KJ, Shteyn V, Bowers SR, Oderberg I, Karpen GH. Assembly of Drosophila centromeric chromatin proteins during mitosis. PLoS Genet. 2011;7(5):e1002068.

53. Dunleavy EM, Beier NL, Gorgescu W, Tang J, Costes SV, Karpen GH. The cell cycle timing of centromeric chromatin assembly in Drosophila meiosis is distinct from mitosis yet requires CAL1 and CENP-C. PLoS Biol. 2012;10(12):e1001460.

54. Lidsky PV, Sprenger F, Lehner CF. Distinct modes of centromere protein dynamics during cell cycle progression in Drosophila S2R+ cells. J Cell Sci. 2013;126(Pt 20):4782–93.

55. Pauleau AL, Bergner A, Kajtez J, Erhardt S. The checkpoint protein Zw10 connects CAL1-dependent CENP-A centromeric loading and mitosis duration in Drosophila cells. PLoS Genet. 2019;15(9):e1008380.

56. Ranjan R, Snedeker J, Chen X. Asymmetric Centromeres Differentially Coordinate with Mitotic Machinery to Ensure Biased Sister Chromatid Segregation in Germline Stem Cells. Cell Stem Cell. 2019;25(5):666–81 e5.

57. Dunleavy EM, Almouzni G, Karpen GH. H3.3 is deposited at centromeres in S phase as a placeholder for newly assembled CENP-A in G(1) phase. Nucleus. 2011;2(2):146–57.

58. Perea-Resa C, Bury L, Cheeseman IM, Blower MD. Cohesin Removal Reprograms Gene Expression upon Mitotic Entry. Mol Cell. 2020;78(1):127–40 e7.

59. Ghosh S, Lehner CF. Incorporation of CENP-A/CID into centromeres during early Drosophila embryogenesis does not require RNA polymerase II-mediated transcription. Chromosoma. 2022;131(1-2):1–17.

60. Raychaudhuri N, Dubruille R, Orsi GA, Bagheri HC, Loppin B, Lehner CF. Transgenerational propagation and quantitative maintenance of paternal centromeres depends on Cid/Cenp-A presence in Drosophila sperm. PLoS Biol. 2012;10(12):e1001434.

61. Hemmer L, Negm S, Geng X, Courret C, Navarro-Domínguez B, Speece I, et al. Centromere-associated retroelement evolution in Drosophila melanogaster reveals an underlying conflict. BioRXiv. 2023.

62. Sullivan BA, Karpen GH. Centromeric chromatin exhibits a histone modification pattern that is distinct from both euchromatin and heterochromatin. Nat Struct Mol Biol. 2004;11(11):1076–83.

63. Jachowicz JW, Bing X, Pontabry J, Boskovic A, Rando OJ, Torres-Padilla ME. LINE-1 activation after fertilization regulates global chromatin accessibility in the early mouse embryo. Nat Genet. 2017;49(10):1502–10.

64. Limouse C, Smith OK, Jukam D, Fryer KA, Greenleaf WJ, Straight AF. Global mapping of RNA-chromatin contacts reveals a proximity-dominated connectivity model for ncRNA-gene interactions. Nat Commun. 2023;14(1):6073.

65. Decker CJ, Burke JM, Mulvaney PK, Parker R. RNA is required for the integrity of multiple nuclear and cytoplasmic membrane-less RNP granules. EMBO J. 2022;41(9):e110137.

66. Rhine K, Vidaurre V, Myong S. RNA Droplets. Annu Rev Biophys. 2020;49:247–65.

67. Schittenhelm RB, Althoff F, Heidmann S, Lehner CF. Detrimental incorporation of excess Cenp-A/Cid and Cenp-C into Drosophila centromeres is prevented by limiting amounts of the bridging factor Cal1. J Cell Sci. 2010;123(Pt 21):3768–79.

68. Pfaffl MW. A new mathematical model for relative quantification in real-time RT-PCR. Nucleic Acids Res. 2001;29(9):e45.

69. Hanlon SL, Miller DE, Eche S, Hawley RS. Origin, Composition, and Structure of the Supernumerary B Chromosome of Drosophila melanogaster. Genetics. 2018;210(4):1197–212.

70. Saunders A, Core LJ, Sutcliffe C, Lis JT, Ashe HL. Extensive polymerase pausing during Drosophila axis patterning enables high-level and pliable transcription. Genes Dev. 2013;27(10):1146–58.

71. Martin M. Cutadapt Removes Adapter Sequences From High-Throughput Sequencing Reads. EMBnet Journal. 2011;Vol 17, No 1.

72. Langmead B, Salzberg SL. Fast gapped-read alignment with Bowtie 2. Nature methods. 2012;9(4):357–9.

73. Quinlan AR, Hall IM. BEDTools: a flexible suite of utilities for comparing genomic features. Bioinformatics. 2010;26(6):841–2.

74. Robinson JT, Thorvaldsdottir H, Winckler W, Guttman M, Lander ES, Getz G, et al. Integrative genomics viewer. Nat Biotechnol. 2011;29(1):24–6.

75. Langmead B, Trapnell C, Pop M, Salzberg SL. Ultrafast and memory-efficient alignment of short DNA sequences to the human genome. Genome Biol. 2009;10(3):R25.

76. Rhie A, Walenz BP, Koren S, Phillippy AM. Merqury: reference-free quality, completeness, and phasing assessment for genome assemblies. Genome Biol. 2020;21(1):245.

77. Buenrostro JD, Wu B, Chang HY, Greenleaf WJ. ATAC-seq: A Method for Assaying Chromatin Accessibility Genome-Wide. Curr Protoc Mol Biol. 2015;109:21 9 1–9 9.

78. Ramirez F, Ryan DP, Gruning B, Bhardwaj V, Kilpert F, Richter AS, et al. deepTools2: a next generation web server for deep-sequencing data analysis. Nucleic Acids Res. 2016;44(W1):W160–5.

