## Supplemental Figures for "Transcription of a centromere-enriched retroelement and local retention of its RNA are significant features of the CENP-A chromatin landscape"

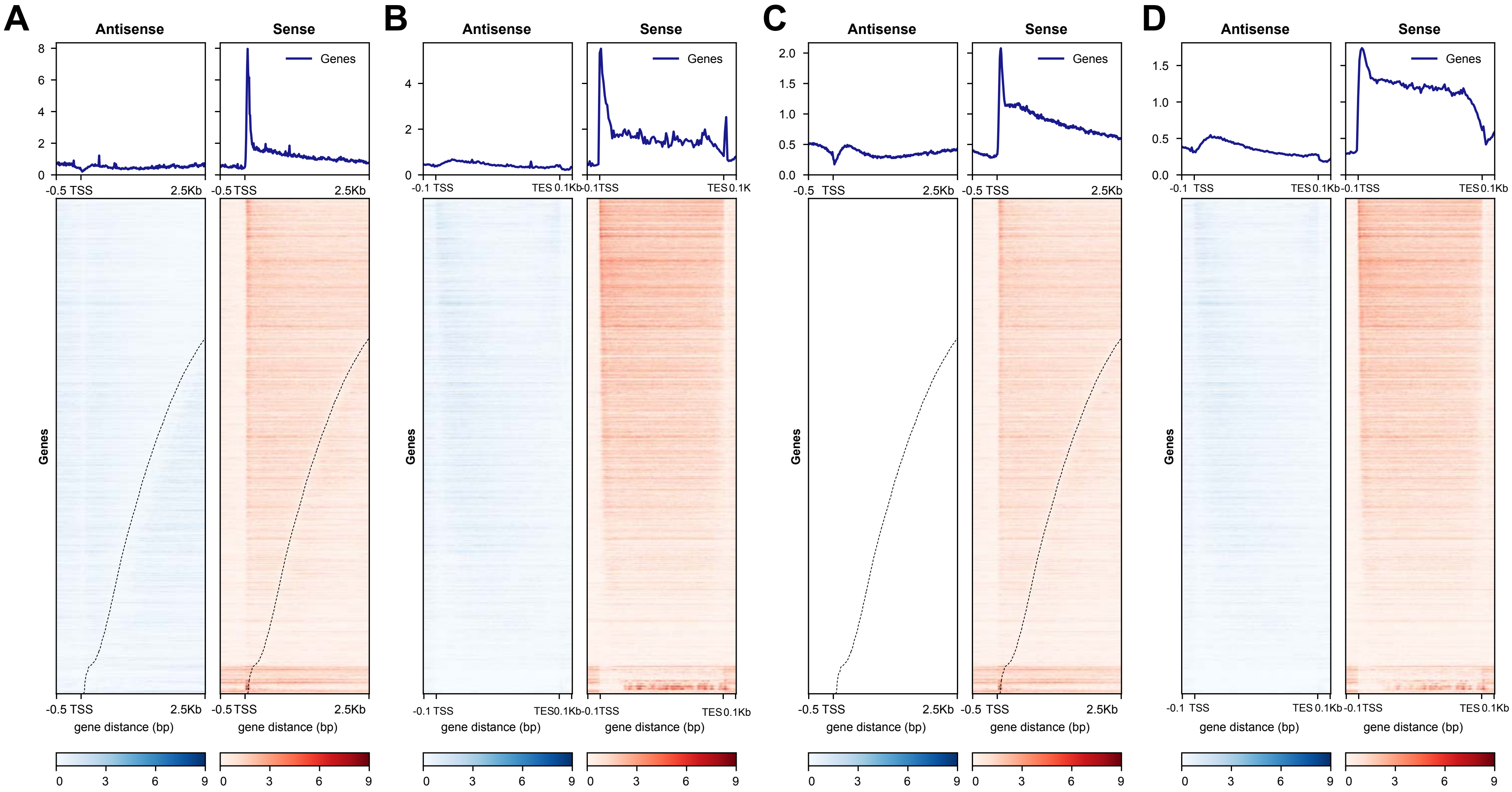
A
B
C
D
Antisense
Sense
Genes
gene distance (bp)
-0.5 TSS
2.5Kb
-0.1 TSS
TES 0.1Kb

Fig.S1

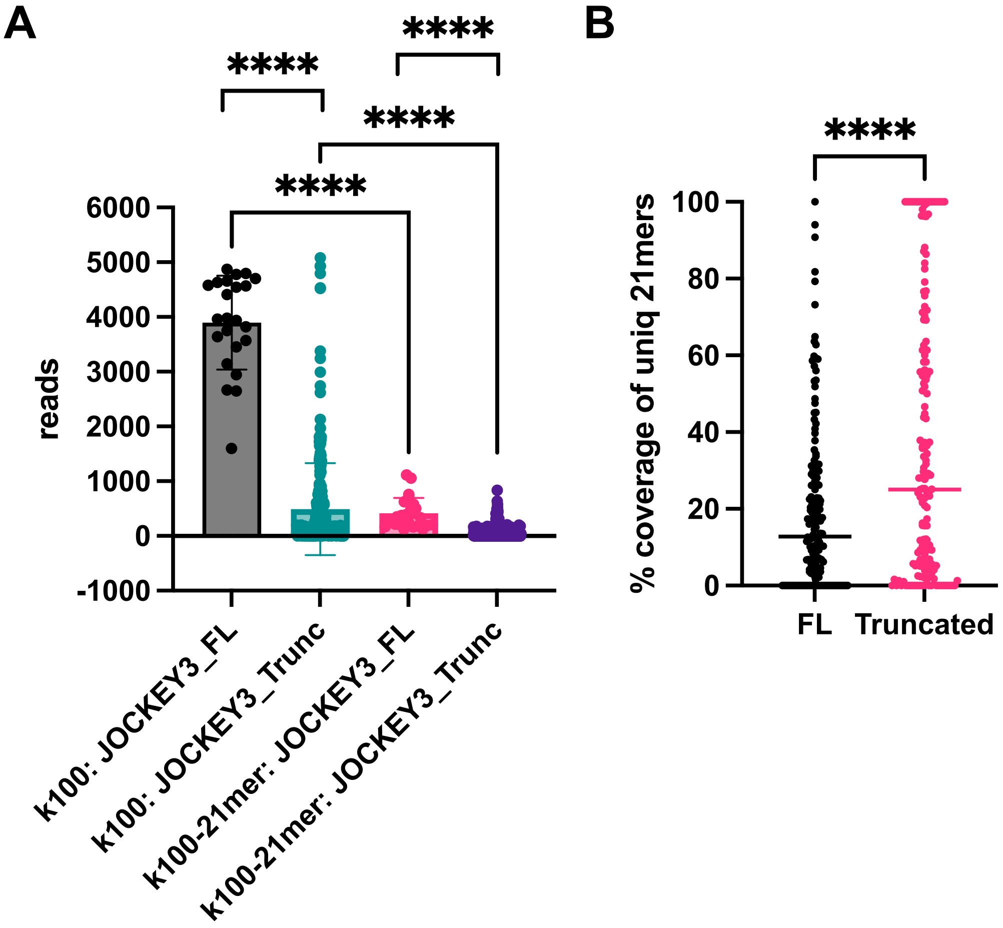
A
B
****
****
****
****
****
reads
6000
5000
4000
3000
2000
1000
0
-1000
k100: JOCKEY3_FL
k100: JOCKEY3_Trunc
k100-21mer: JOCKEY3_FL
k100-21mer: JOCKEY3_Trunc
% coverage of uniq 21mers
100
80
60
40
20
0
FL
Truncated

Fig. S2

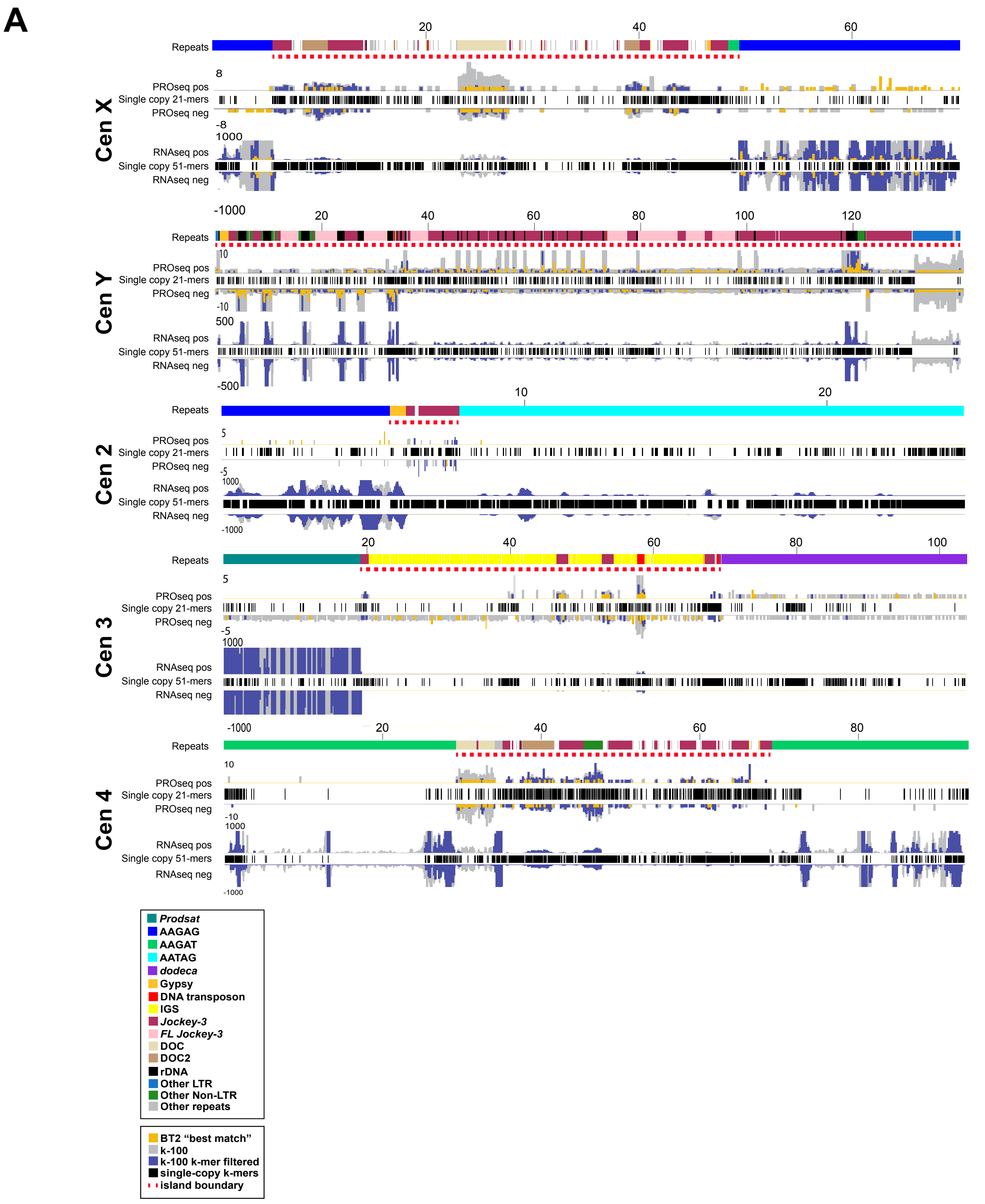

A
Cen X
Repeats
PROseq pos
Single copy 21-mers
PROseq neg
RNAseq pos
Single copy 51-mers
RNAseq neg
Cen Y
Cen 2
Cen 3
Cen 4
Prodsat
AAGAG
AAGAT
AATAG
dodeca
Gypsy
DNA transposon
IGS
Jockey-3
FL Jockey-3
DOC
DOC2
rDNA
Other LTR
Other Non-LTR
Other repeats
BT2 "best match"
k-100
k-100 k-mer filtered
single-copy k-mers
island boundary


Fig. S3

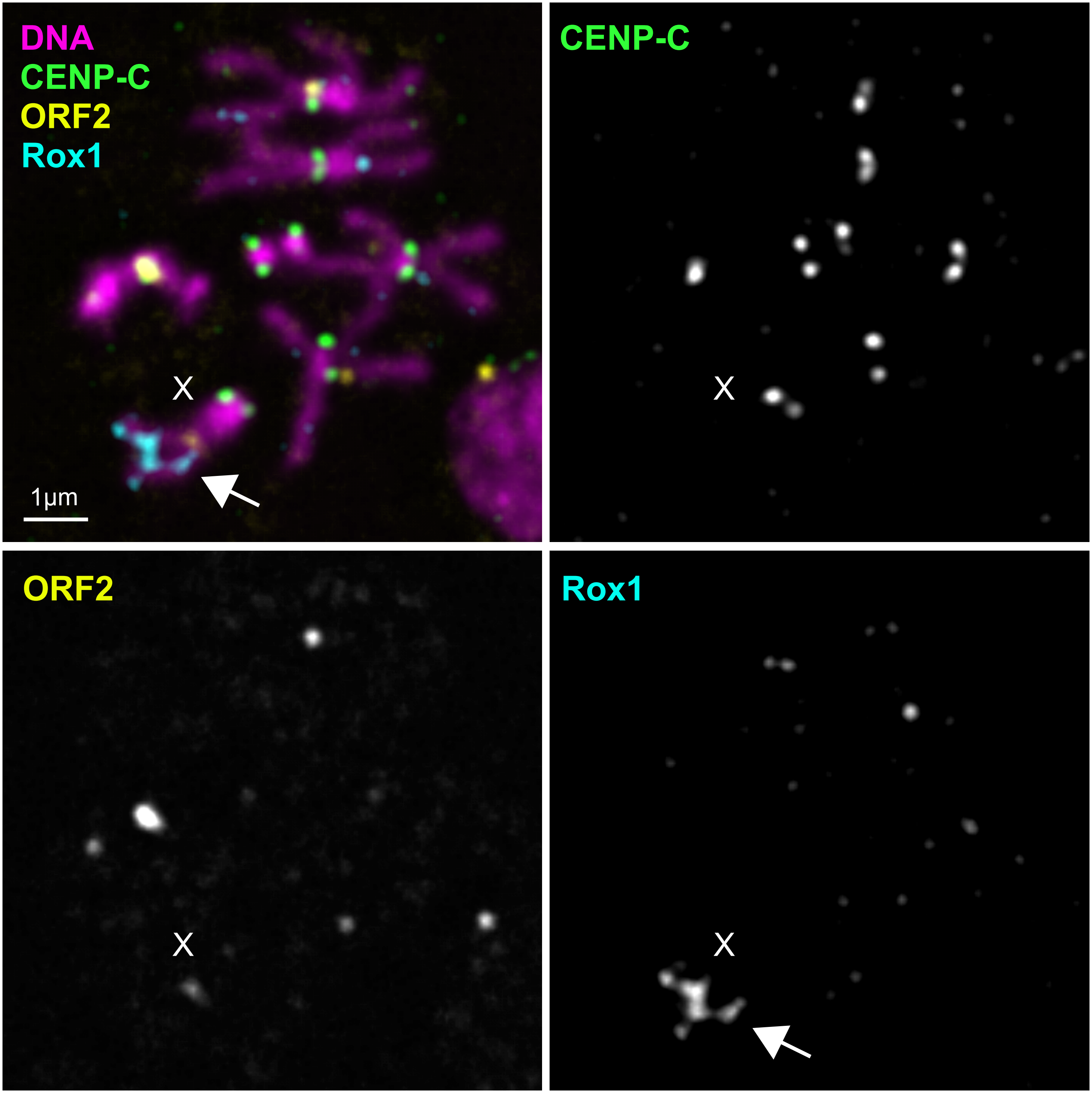


Fig. S4

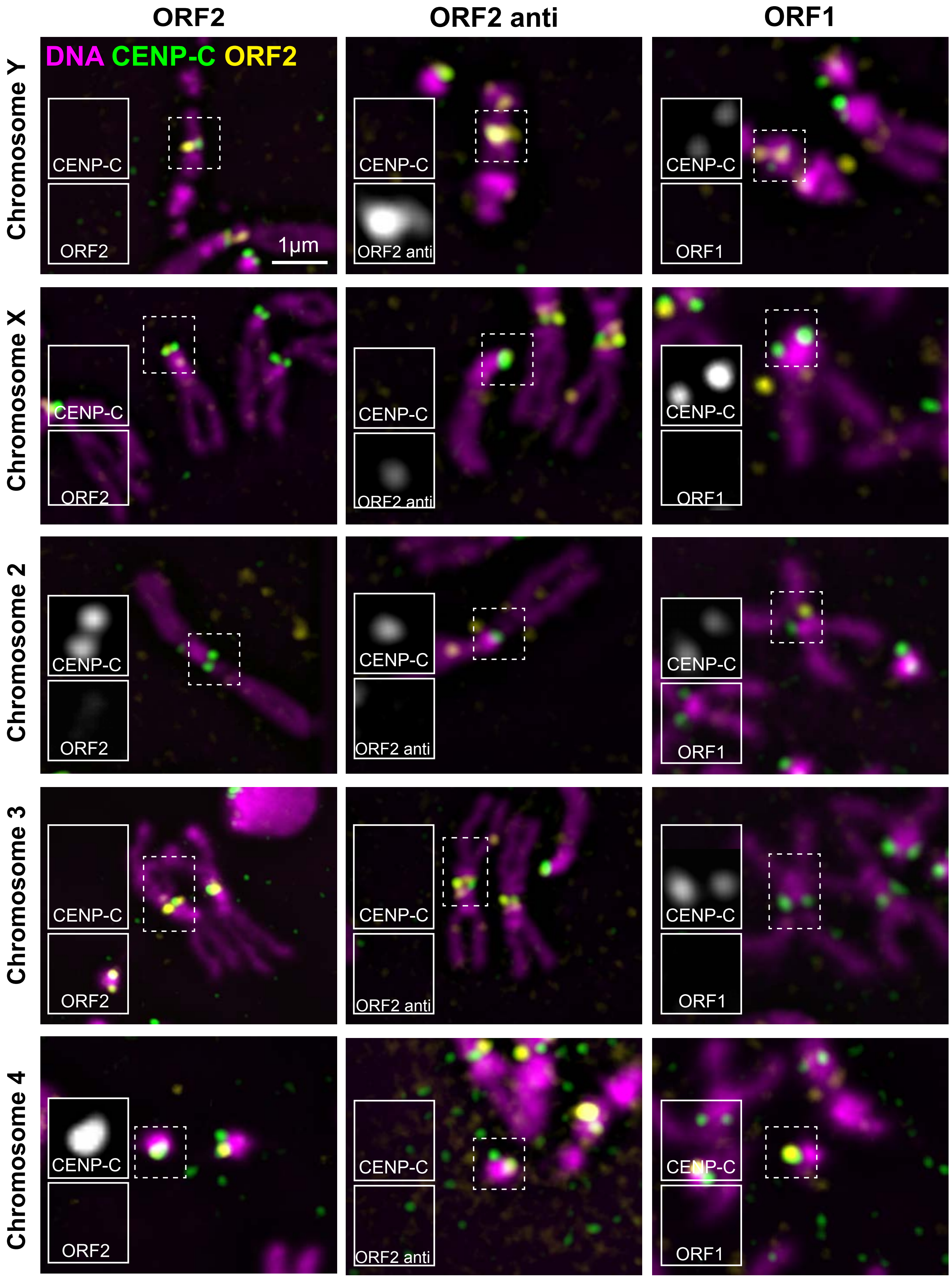

ORF2
ORF2 anti
ORF1
DNA CENP-C ORF2
Chromosome Y
Chromosome X
Chromosome 2
Chromosome 3
Chromosome 4
CENP-C
ORF2
ORF2 anti
ORF1
1μm


### Fig. S5

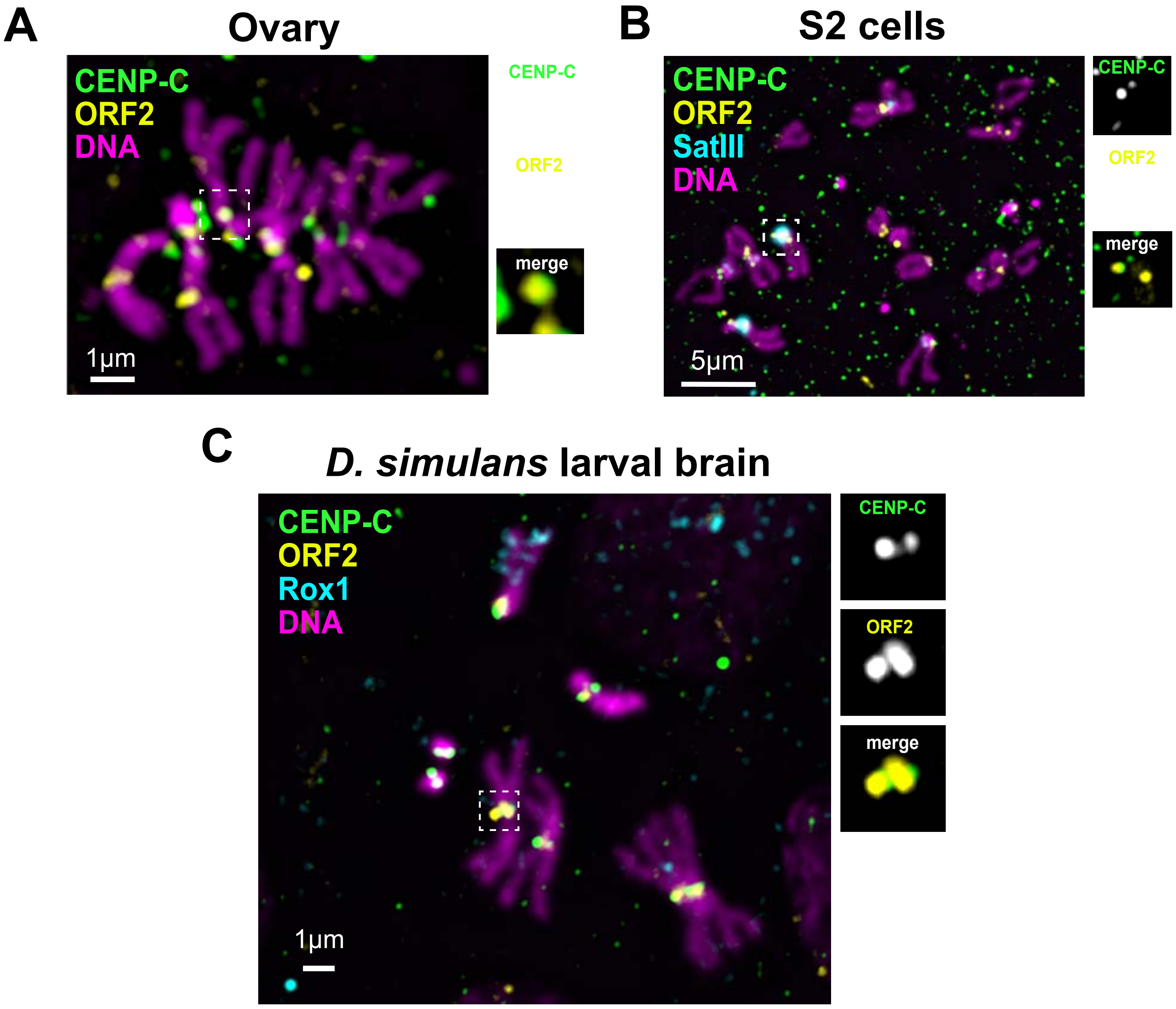
A
Ovary
CENP-C
ORF2
DNA
1µm
CENP-C
ORF2
merge
B
S2 cells
CENP-C
ORF2
SatIII
DNA
5µm
CENP-C
ORF2
merge
C
D. simulans larval brain
CENP-C
ORF2
Rox1
DNA
1µm
CENP-C
ORF2
merge

Fig. S6

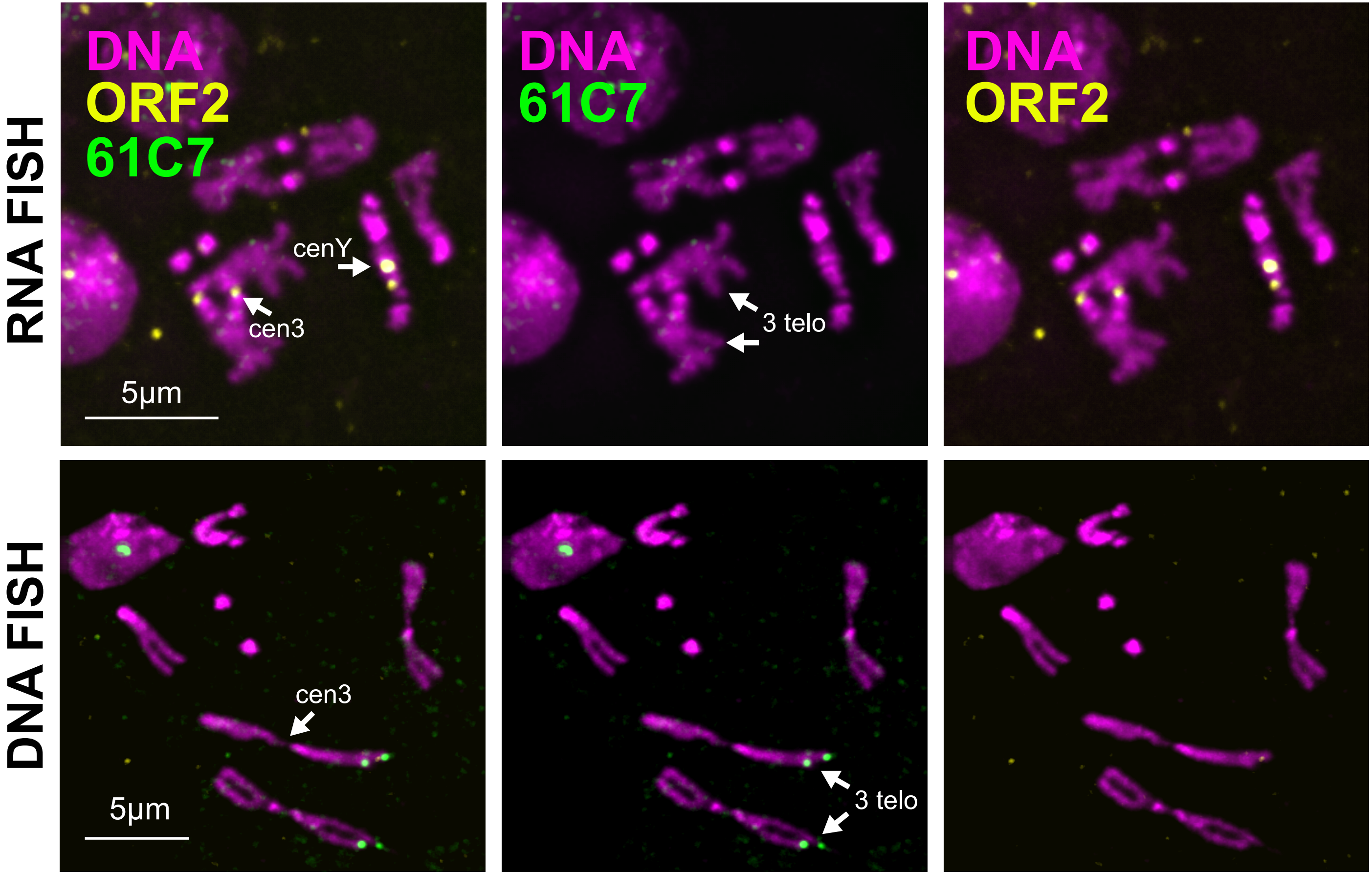

RNA FISH
DNA FISH
DNA
ORF2
61C7
cenY
cen3
5µm
DNA
61C7
3 telo
DNA
ORF2
cen3
5µm
3 telo


Fig. S7

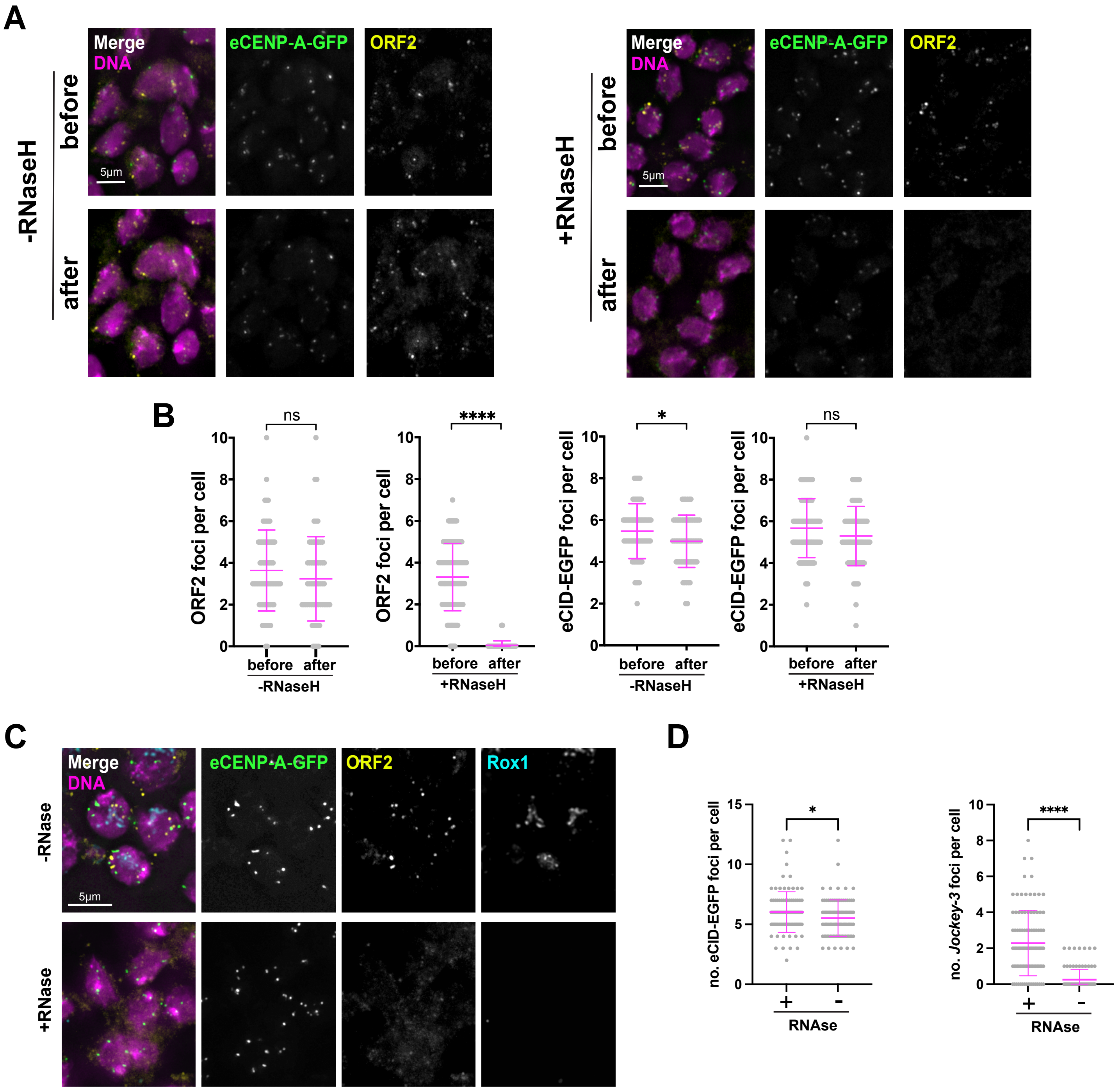
A
Merge
DNA
eCENP-A-GFP
ORF2
-RNaseH
before
after
5µm
+RNaseH
B
ns
****
*
ORF2 foci per cell
eCID-EGFP foci per cell
before
after
-RNaseH
+RNaseH
C
Rox1
-RNase
+RNase
D
no. eCID-EGFP foci per cell
no. Jockey-3 foci per cell
RNAse
+
-

Fig. S8

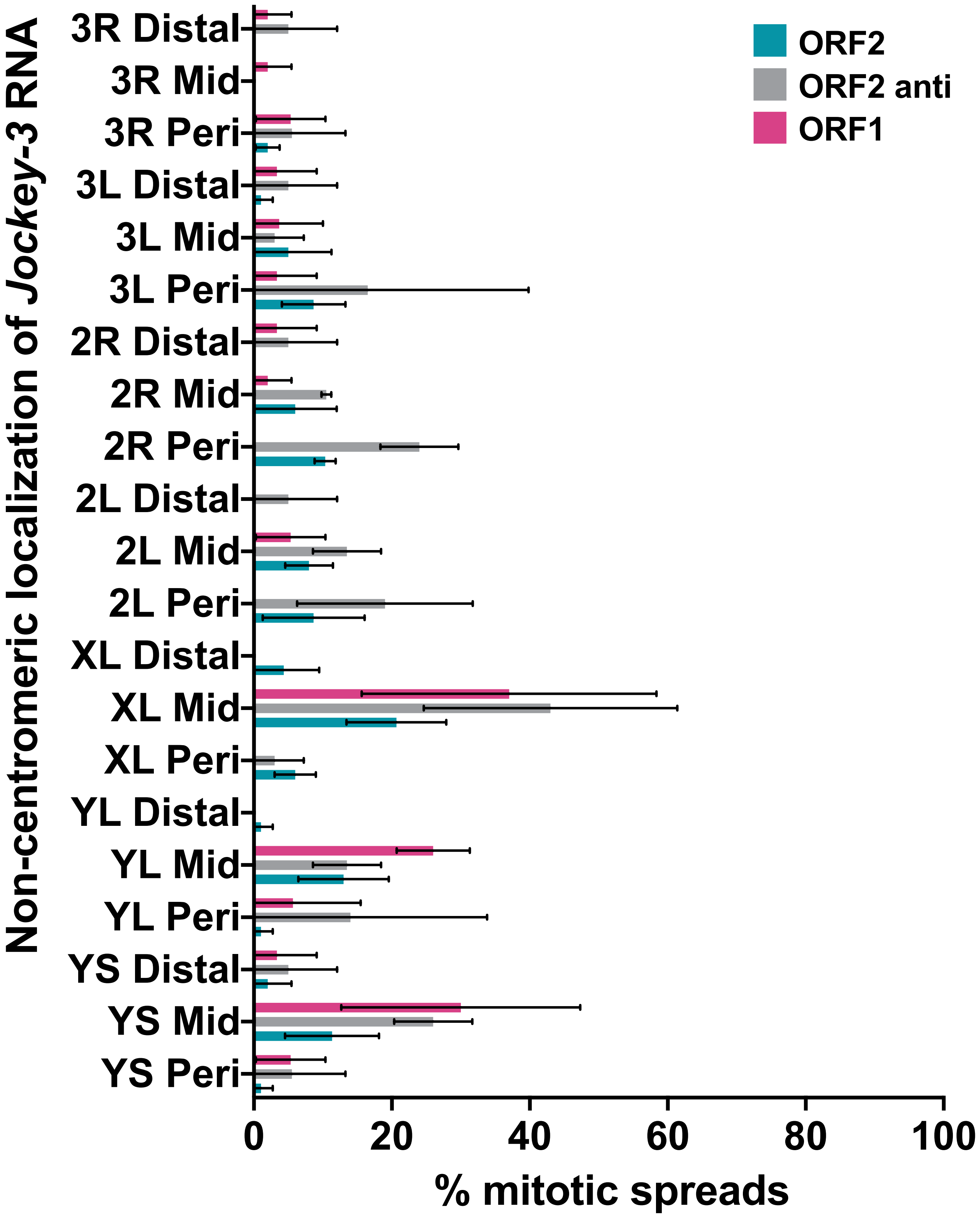

Non-centromeric localization of Jockey-3 RNA
3R Distal
3R Mid
3R Peri
3L Distal
3L Mid
3L Peri
2R Distal
2R Mid
2R Peri
2L Distal
2L Mid
2L Peri
XL Distal
XL Mid
XL Peri
YL Distal
YL Mid
YL Peri
YS Distal
YS Mid
YS Peri
0
20
40
60
80
100
% mitotic spreads
ORF2
ORF2 anti
ORF1


Fig. S9

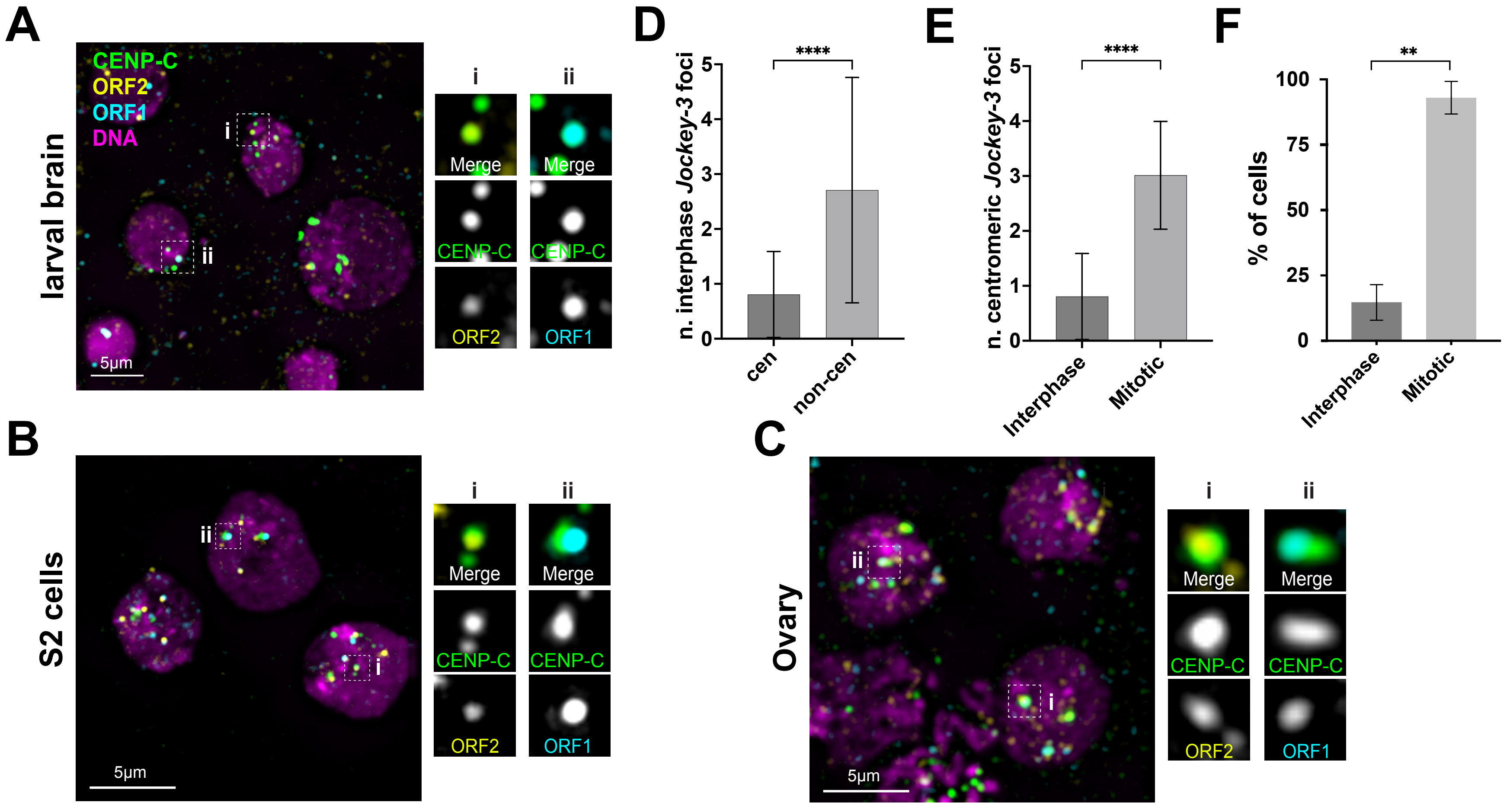
A
CENP-C
ORF2
ORF1
DNA
larval brain
i
ii
Merge
CENP-C
ORF2
ORF1
5µm
B
S2 cells
C
Ovary
D
n. interphase Jockey-3 foci
****
cen
non-cen
E
n. centromeric Jockey-3 foci
****
Interphase
Mitotic
F
% of cells
**
0
25
50
75
100

Fig. S10

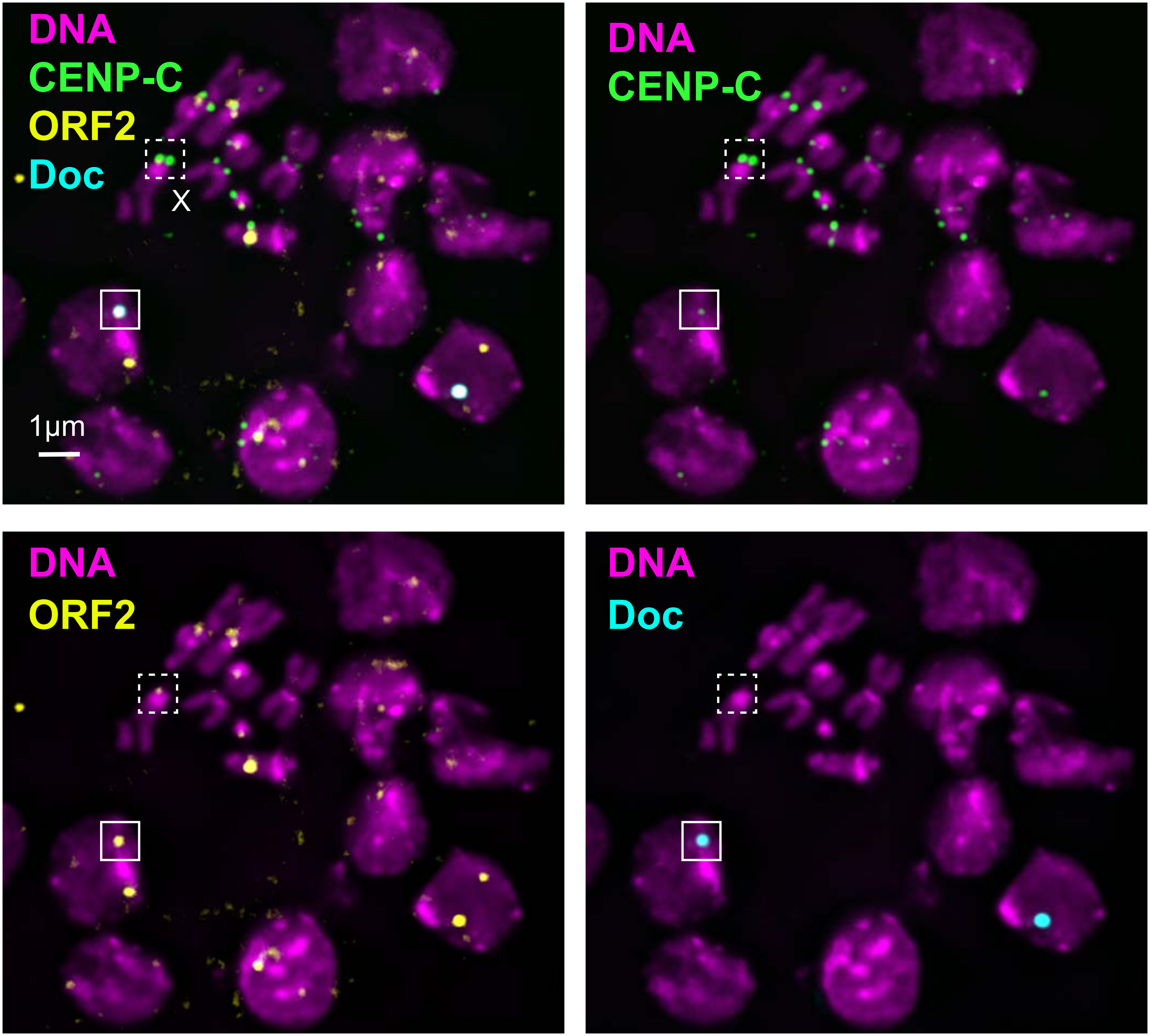


**Fig. S11**

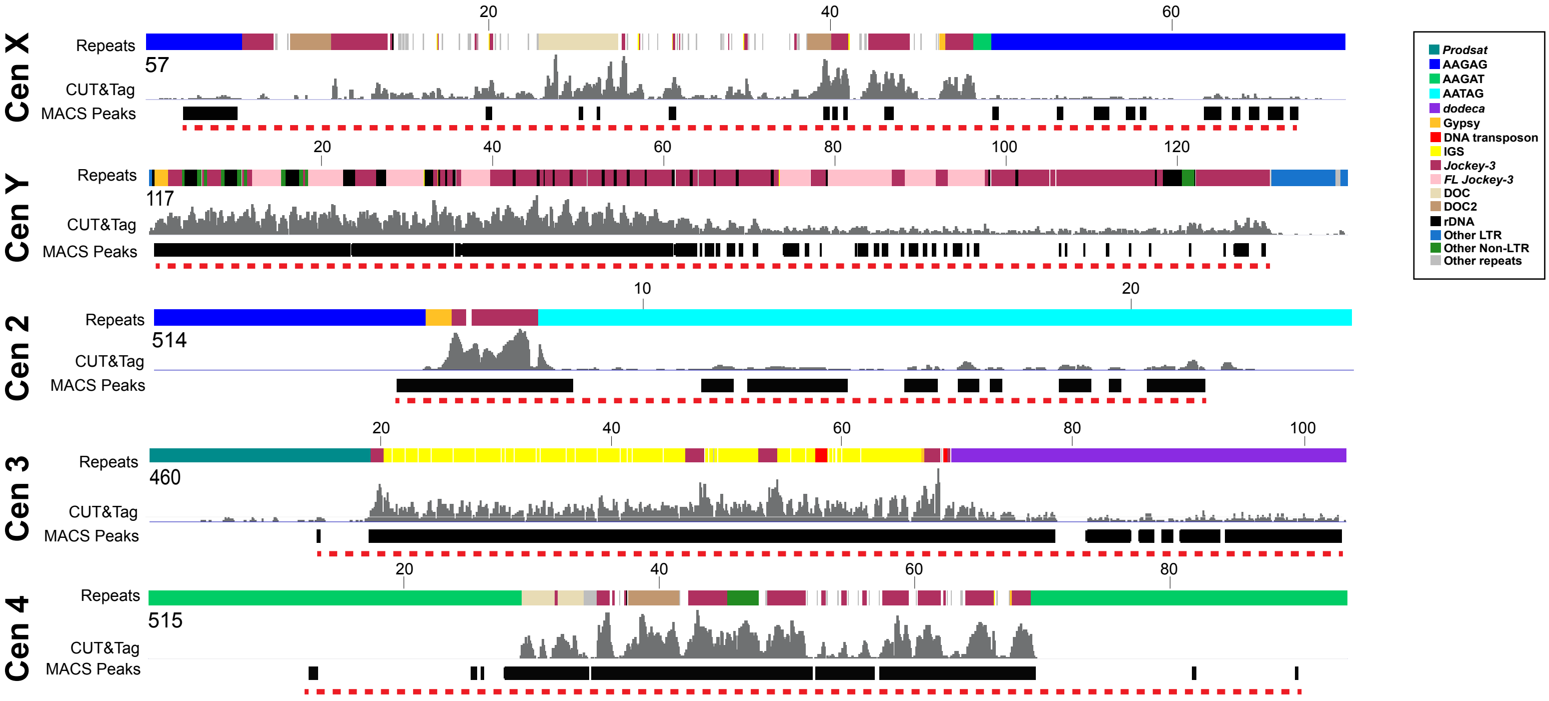
Cen X
Repeats
CUT&Tag
MACS Peaks
57
20
40
60
Cen Y
117
100
120
Cen 2
514
10
Cen 3
460
80
Cen 4
515
Prodsat
AAGAG
AAGAT
AATAG
dodeca
Gypsy
DNA transposon
IGS
Jockey-3
FL Jockey-3
DOC
DOC2
rDNA
Other LTR
Other Non-LTR
Other repeats

**Fig. S12**

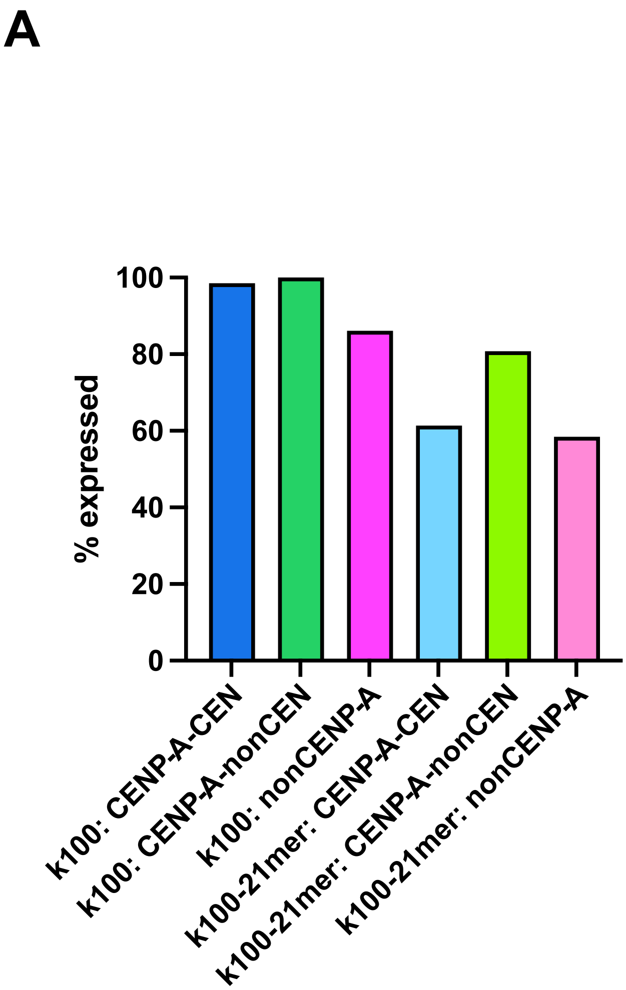
A
% expressed
100
80
60
40
20
0
k100: CENP-A-CEN
k100: CENP-A-nonCEN
k100: nonCENP-A
k100-21mer: CENP-A-CEN
k100-21mer: CENP-A-nonCEN
k100-21mer: nonCENP-A

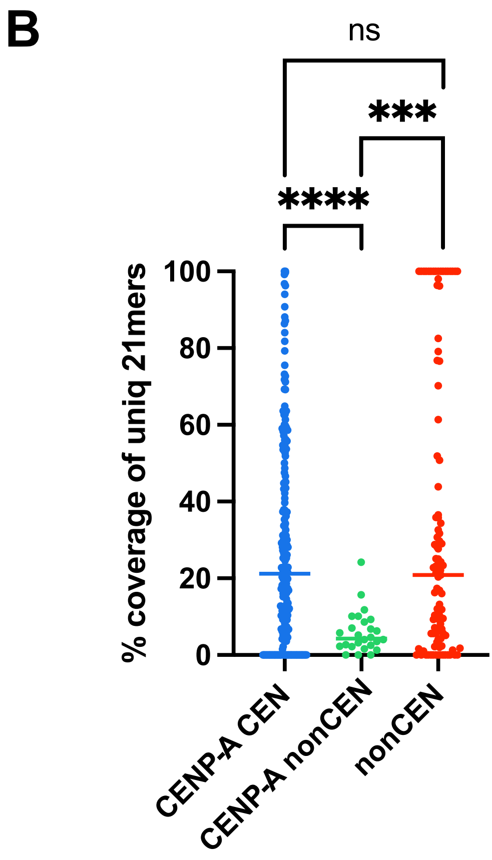
B
ns
***
****
% coverage of uniq 21mers
100
80
60
40
20
0
CENP-A CEN
CENP-A nonCEN
nonCEN

Fig. S13
